# A Buprenorphine depot formulation provides effective sustained post-surgical analgesia for 72h in mouse femoral fracture models

**DOI:** 10.1101/2022.07.05.498859

**Authors:** Angelique Wolter, Christian H. Bucher, Sebastian Kurmies, Viktoria Schreiner, Frank Konietschke, Katharina Hohlbaum, Robert Klopfleisch, Max Löhning, Christa Thöne-Reineke, Frank Buttgereit, Jörg Huwyler, Paulin Jirkof, Anna E. Rapp, Annemarie Lang

**Affiliations:** Charité – Universitätsmedizin Berlin, corporate member of Freie Universität Berlin and Humboldt-Universität zu Berlin, Department of Rheumatology and Clinical Immunology, Berlin, Germany; German Rheumatism Research Centre (DRFZ), a Leibniz Institute, Berlin, Germany; Institute of Animal Welfare, Animal Behavior and Laboratory Animal Science, Department of Veterinary Medicine, Freie Universität Berlin, Berlin, Germany; Julius Wolff Institute, Charité – Universitätsmedizin Berlin, Berlin, Germany; Berlin Institute of Health Center for Regenerative Therapies (BCRT), Charité – Universitätsmedizin Berlin, Berlin, Germany; Division of Pharmaceutical Technology, Department of Pharmaceutical Sciences, University of Basel, Basel, Switzerland; Charité-Universitätsmedizin Berlin, Corporate Member of Freie Universität Berlin and Humboldt-Universität zu Berlin, Institute of Biometry and Clinical Epidemiology, Berlin, Germany; German Federal Institute for Risk Assessment (BfR), German Centre for the Protection of Laboratory Animals (Bf3R), Berlin, Germany; Institute of Veterinary Pathology, Department of Veterinary Medicine, Freie Universität Berlin, Berlin, Germany; Center for Surgical Research, University Hospital Zurich, University Zurich, Zurich, Switzerland; Office for Animal Welfare and 3Rs, University of Zurich, Zurich, Switzerland; Dr. Rolf M. Schwiete Research Unit for Osteoarthritis, Department of Orthopedics (Friedrichsheim), University Hospital Frankfurt, Goethe University, Frankfurt, Germany; Departments of Orthopaedic Surgery and Bioengineering, University of Pennsylvania, PA, United States

**Author notes:** these authors contributed equally (Anna E. Rapp, Annemarie Lang). Correspondences: Angelique Wolter, Institute of Animal Welfare, Animal Behavior and Laboratory Animal Science, Department of Veterinary Medicine, Freie Universität Berlin, Berlin, Germany, Annemarie Lang, Departments of Orthopaedic Surgery and Bioengineering, University of Pennsylvania, PA, United States.

## Abstract

Adequate pain management is essential for ethical and scientific reasons in animal experiments and should completely cover the period of expected pain without the need for frequent re-application. However, current depot formulations of Buprenorphine are only available in the USA and have limited duration of action. Recently, a new microparticulate Buprenorphine formulation (BUP-Depot) for sustained release has been developed as an alternative product within Europe. Pharmacokinetics indicate a potential effectiveness for about 72h. Here, we investigated whether the administration of the BUP-Depot ensures continuous and sufficient analgesia in two mouse femoral fracture models and could therefore serve as a potent alternative to the application of Tramadol via drinking water. Both protocols were examined for analgesic effectiveness, side effects on experimental readout, and effects on fracture healing outcomes in male and female C57BL/6N mice. The BUP-Depot provided effective analgesia for 72h, comparable to the effectiveness of Tramadol in drinking water. Fracture healing outcome was not different between analgesic regimes. The availability of a Buprenorphine depot formulation for laboratory animals in Europe would be a beneficial addition for extended pain relief in mice, thereby increasing animal welfare.

## Introduction

Animals – especially mice – are still widely used and required in fundamental and translational research to study the complexity of biological and pathophysiological processes. Therefore, the active implementation of the 3R principle (Replace – Reduce – Refine), with a particular importance of *Refinement* forms the indispensable basis for a humane approach to conduct animal experiments. Thus, adequate pain assessment and medication in animals before, during and after the experimental procedure are crucial to decrease suffering and ensure data quality. Insufficiently treated pain and handling-induced stress can affect animal behavior and physiological responses, especially in an immunological context, leading to potential bias in the scientific outcomes and reduced reproducibility ^1–6^. However, evidence-based data on individual pain management efficiencies in surgical mouse models is still rare ^6,7^ and the reporting quality of the used analgesic protocols is often insufficient ^8,9^.

After surgical intervention, the potent opioid Buprenorphine is often used for pain relieve in rodents ^10,11^. Due to a short half-life of approximately 3 hours ^12–14^, frequent injections are required, resulting in repeated handling of the animals. However, the commonly reported application intervals of Buprenorphine of every 8 – 12h can lead to pain peaks due to insufficient analgesic coverage ^9,15^. The parenteral application of other opioids such as Morphine, Tramadol and Fentanyl is also rather not suitable for pain alleviation in rodents, as their half-life is even shorter ^13,16,17^.

To reduce handling-associated stress and to ensure continuous analgesic coverage, an alternative application route in form of administration of Buprenorphine or Tramadol with the drinking water has been routinely used e.g., in orthopedics-related mouse models ^18–21^. However, as the uptake of analgesics with the drinking water is dependent on the drinking frequency and intake amount, the overall effectiveness of this treatment strategy might be highly influenced by e.g., reduced activity and water intake after anesthesia/surgery and circadian activity ^22^. A drug formulation that extends the analgesic effect due to sustained parenteral drug release could overcome such challenges and serve as a powerful tool to further refine today’s analgesic regimens in animal experiments. However, current depot/sustained-release formulations of Buprenorphine for mice and rats are either only available for dedicated research purposes or are only available in the United States of America (USA), e.g., Buprenorphine SR-LAB (ZooPharm) or Ethiqa XR (Fidelis Animal Health). Attempts to import these products to Europe have failed due to missing approval through the European Medical Evaluation Agency (EMEA).

To fill the gap, Schreiner et al. successfully developed a poly-lactic-co-glycolic acid (PLGA) based microparticulate drug formulation for sustained drug release of buprenorphine ^23,24^. In a proof-of-concept study, they observed therapeutic-relevant drug levels of the sustained-release Buprenorphine (BUP-Depot) in the brain for more than 24h, an antinociceptive effect in the hot plate test, and pain relief after a minor abdominal surgery in female C57BL/6J mice for at least 72h ^23^.

This present study therefore aims at exploring the analgesic capacities of the newly developed BUP-Depot compared to Tramadol in the drinking water and its potential to improve animal welfare in a wider range of mouse models in Europe. To test the effectiveness of the newly developed BUP-Depot in a preclinical setting of surgical interventions, we here compared the analgesic capacities of the BUP-Depot to the established application of Tramadol with the drinking water. Both pain management protocols were examined for their analgesic efficacy and adverse effects on experimental readouts in two femoral fracture models using rigid and flexible external fixators. To consider potential sex-dependent differences in response to the analgesic protocol, male and female mice were included. We monitored (i) general parameters of wellbeing e.g., body weight, food and water intake, nestbuilding and explorative behavior, composite score, and (ii) model-specific pain parameters including walking behavior (limp score) and CatWalk analysis. In addition, fracture healing outcomes were examined at the end of the study to exclude negative influences on the regeneration process.

## Results

To investigate the analgesic efficacy of BUP-Depot, we chose an integrative study design to i) generate intra-individual controls for the behavioral assessments and ii) reduce animal numbers used in this study (**Fig. 1**). In brief, animals underwent a first intervention consisting of isoflurane anesthesia and administration of the assigned analgesics. Assessments were performed at 12h, 24h, 48h and 72h (referred to in the following as “post-anesthesia”). 14 days after the first intervention, the same animals underwent a second intervention including isoflurane anesthesia, administration of the respective analgesics and an additional osteotomy on the left femur. The same assessments as post-anesthesia were performed at 12h, 24h, 48h and 72h (in the following referred to as “post-osteotomy”) (**Fig 1b**). Of note, mice did not undergo osteotomy during the first intervention (post-anesthesia), but were already assigned to the respective fixation, leading to group descriptions of “rigid fixation” or “flexible fixation” even after anesthesia.

**Figure 1.**
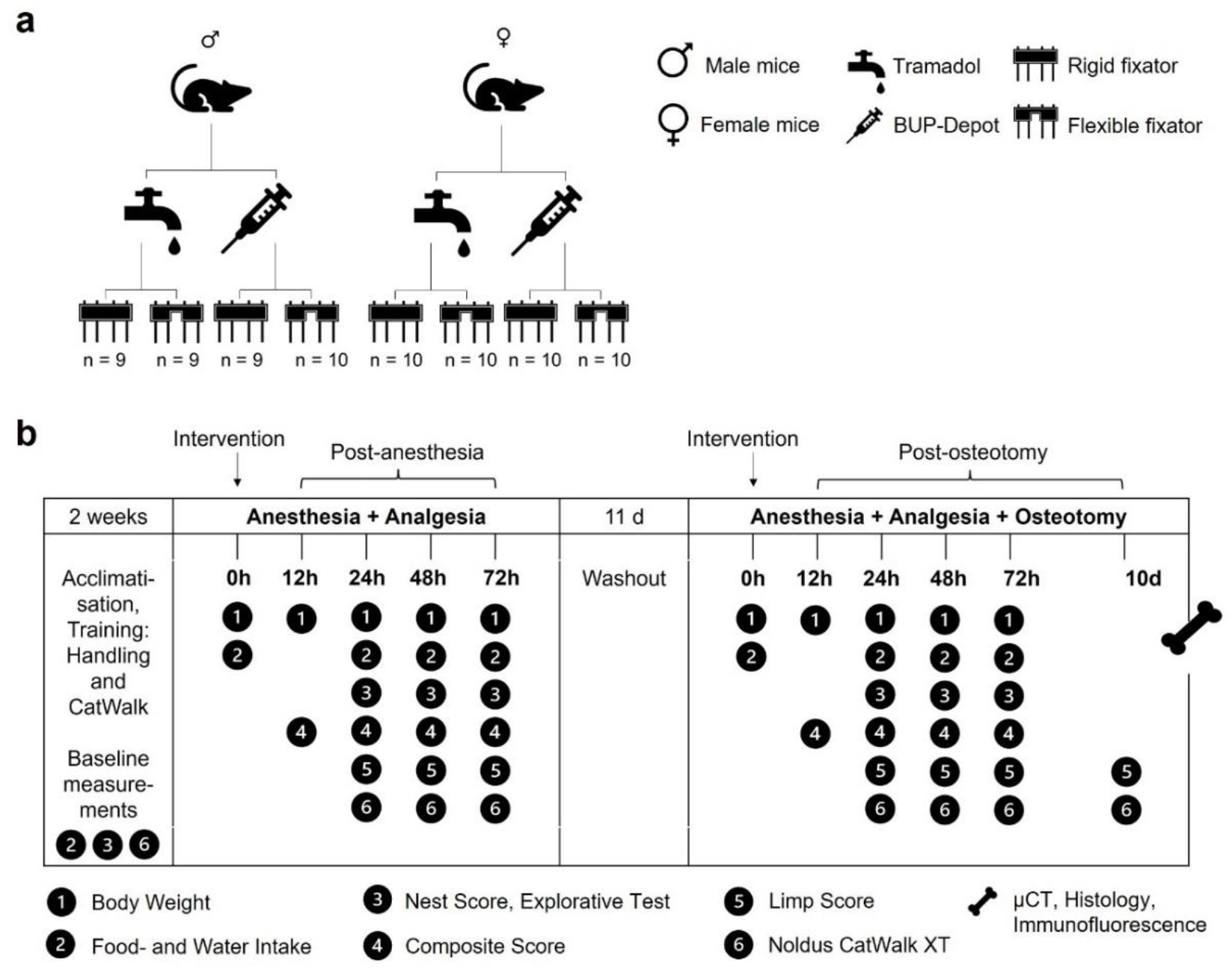
Group assignment and study design. Overview on (**a**) the group assignment, and (**b**) time points and measurements of the different parameters. To assure a rapid onset of the analgesic effect, each animal received a single s.c. dose of Temgesic (1 mg/kg) at the beginning of each intervention. Depending on the assigned analgesic protocol, mice additionally received Tramadol (0.1 mg/ml) in drinking water (provided one day before and for three consecutive days after the interventions) or a single dose of sustained-release BUP-Depot (1.2 mg/kg s.c) was administered at the end of the two interventions.

### Body weight, food and water intake are affected post-anesthesia and post-osteotomy independent of analgesic regime and fixation

To assess general indications for wellbeing, the body weight was monitored at 12h, 24h, 48h and 72h post-anesthesia (initial body weight at 0h – males: 25.96 ± 2.1 g; females: 21.41 ± 1.3 g) and post-osteotomy (initial body weight at 0h – males: 27.61 ± 1.9 g; females: 22.75 ± 1.3 g). The body weight showed a statistically significant reduction in the range of 5% in all groups at 12h and 24h regardless of intervention, sex, fixation, and analgesic regime (**Fig. 2**). At 48h post-anesthesia, body weight normalized in all groups and even exceeded the pre-intervention weight (**Fig. 2a, b**). After osteotomy, we found that male mice showed prolonged body weight loss over 48h, when compared to post-anesthesia (**Fig. 2a**). Female mice had less body weight loss at 24h post-osteotomy compared to post-anesthesia, but a similar recovery at 48h (**Fig. 2b**). The body weight increase at 72h compared to the initial value was higher post-anesthesia than post-osteotomy in all groups independent of sex, analgesia, and fixation. Until osteotomy/euthanasia, body weight was measured every other day with comparable weight development at any time point (**Fig. S1**).

**Figure 2.**
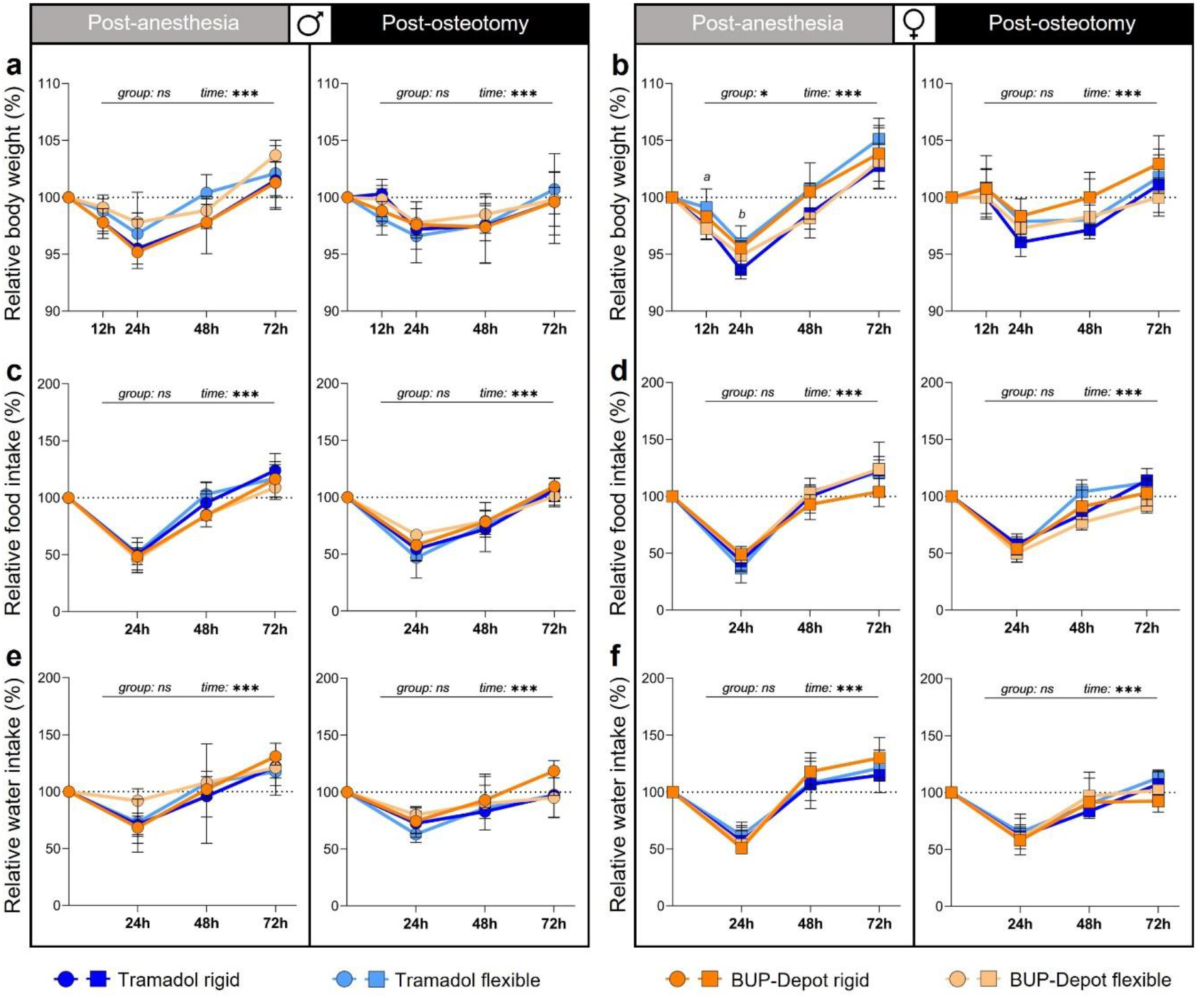
Reduction of body weight as well as food and water intake can be observed at 24h and 48h post-anesthesia and post-osteotomy. (**a, b**) Body weight was measured at 12h, 24h, 48h and 72h; (**c-f**) and food/water intake was measured at 12h, 24h, 48h and 72h post-anesthesia and post-osteotomy. Body weight and food/water intake were normalized to the initial value (0h = 100%). Of note, mice did not undergo osteotomy during the first intervention (= post-anesthesia). However, they were already assigned to their respective groups post-osteotomy. All graphs show median with interquartile range for n = 9–10 (body weight) and n = 4–5 (food/water intake). Non-parametric ANOVA-type test - main effects of time and of group are represented in the graphs; exact *p-values* are listed in Table S1–S3; \**p < 0.05, ***p < 0.001*. To determine group differences Kruskal-Wallis test and Dunn’s posthoc test with Bonferroni correction were performed. *a* – significant difference Tramadol flexible vs. BUP-Depot flexible; *b* – significant difference Tramadol rigid vs. Tramadol flexible.

Food and water intake per cage were assessed at 24h, 48h and 72h post-anesthesia and post-osteotomy. Initial values at 0h covering the previous 24h per cage were as follows: post-anesthesia - males 8.1 ± 0.7 g (food) and 9.2 ± 1.5 ml (water); females 7.8 ± 1.2 g (food) and 9.2 ± 1.2 ml (water); post-osteotomy – males 8.7 ± 1.0 g (food) and 9.9 ± 1.5 ml (water); females 8.4 ± 0.9 g (food) and 9.5 ± 0.9 ml (water). The lowest food and water intake (approximately 50% reduction to initial values) across all groups and sexes was measured 24h after each intervention and reached the level of the initial values at 48h (post-anesthesia) or 72h (post-osteotomy) (**Fig. 2c–f**). No significant main effects were detected between treatment groups (**Table S2, S3**). To rule out any constipating adverse effects of the BUP-Depot, all groups were closely monitored for defecation during the assessments, as constipation is a known-side effect of chronic-opioid usage ^15,25^. However, a reduction in defecation was only noticeable at 12h after both interventions but showed no differences between the Tramadol and BUP-Depot groups. The reduction in defecation was considerably more pronounced in female than in male mice (**Fig. S2**).

### Nest building and explorative behaviors are not influenced by the analgesic regime

To detect model independent changes in spontaneous behavior, monitoring of nest complexity scores and explorative behaviors was performed at 24h, 48h and 72h post-anesthesia and post-osteotomy. Analyses showed no differences in the nest building performance between the Tramadol and BUP-Depot groups post-anesthesia and post-osteotomy as medians ranged between scores of 4.5 to 5 (**Fig. 3**). Explorative behavior was present in all cages with male mice post-anesthesia while reduced in some cages at 24h post-osteotomy (8/20) independent of analgesic regime or fixation. In the female mice, 3 out of 20 cages (Tramadol flexible and BUP-Depot flexible) showed no exploration 24h after both interventions (**Fig. S3**).

**Figure 3.**
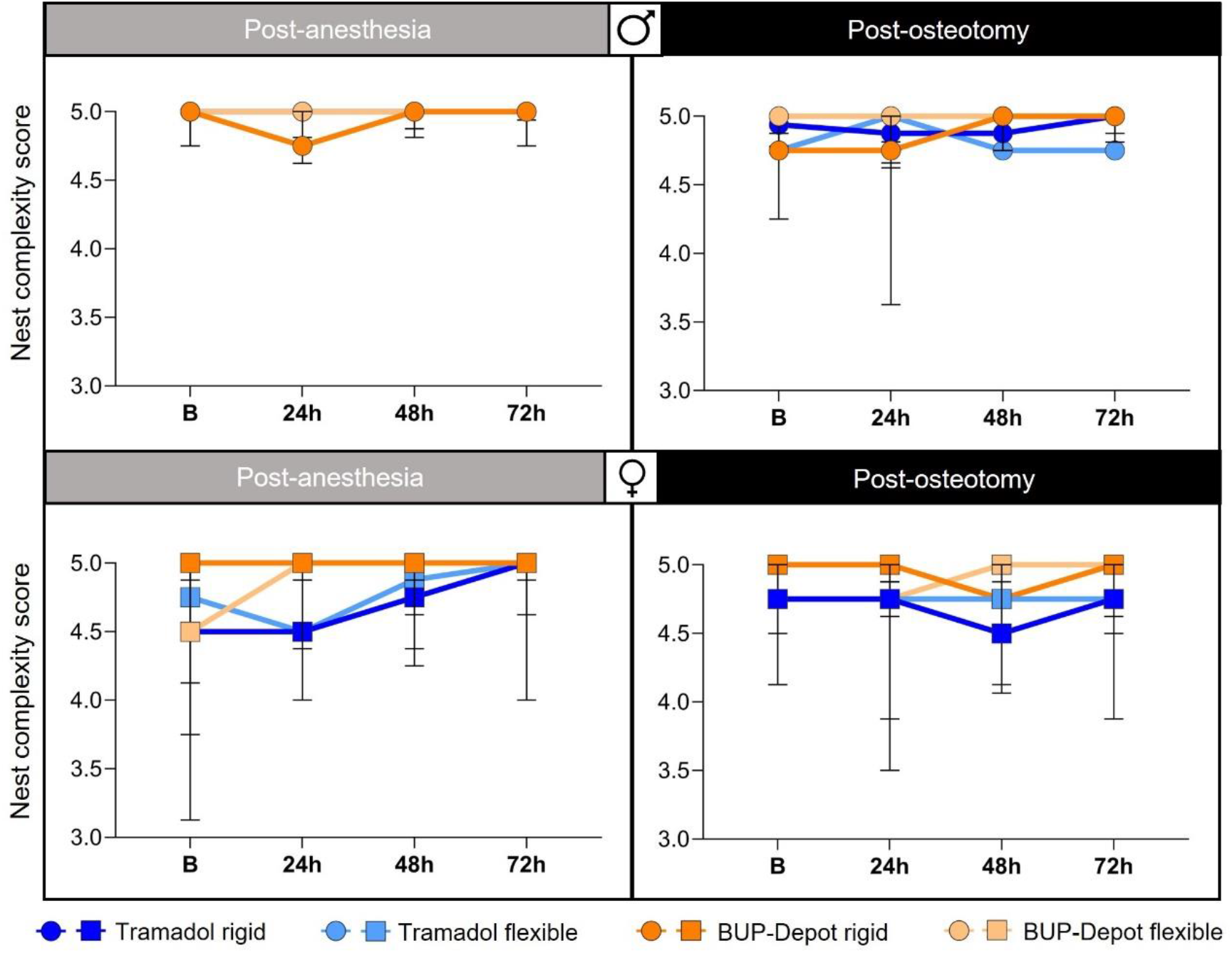
Nest building behavior remains largely unaffected post-anesthesia and post-osteotomy. Nest building was monitored per cage at 24h, 48h and 72h post-anesthesia and post-osteotomy. All graphs show median with interquartile range for n = 4–5 based on cages (pair housing).

### Delta composite pain score indicates limited analgesic capacity of Tramadol in male mice with flexible fixation

The composite pain score combines parameters of facial expression (mouse grimace scale) and overall appearance and was assessed at 24h, 48h and 72h post-anesthesia and post-osteotomy. Since we observed a pronounced influence of the anesthesia and analgesic regime alone (**Fig. S4**; **Table S4–S4.3**), we corrected the individual scores post-osteotomy for the respective scores post-anesthesia to obtain a delta composite pain score, that depicts the isolated effect of the osteotomy without the interfering effects of anesthesia and analgesia. The delta composite pain score was highest in all groups at 12h and 24h after osteotomy, and declined after 48h and 72h, reaching lowest scores in all groups after 72h (significant main time effect in both sexes, *p<0.001;* **Fig. 4a**; **Table S5**). Female mice showed comparable score developments between treatment and fixation groups with the highest median scores (1-1.5) at 12h. In male mice, we found a significantly higher delta composite pain score in the flexible fixation group treated with Tramadol at 24h (median score = 2) and 48h (median score = 1.5) (Kruskal-Wallis test of all groups at 24h: *p= 0.031* and at 48h: *p= <0.001;* **Table S5.1**) when compared to the other groups (Dunn’s posthoc test for Tramadol flexible vs. BUP-Depot flexible, BUP-Depot rigid and Tramadol rigid, respectively: 24h *p= 0.077, 0.042, 0.032;* 48h *p= 0.004, <0.001, 0.018*; **Fig 4a**; **Table S5.2–S5.3**).

**Figure 4.**
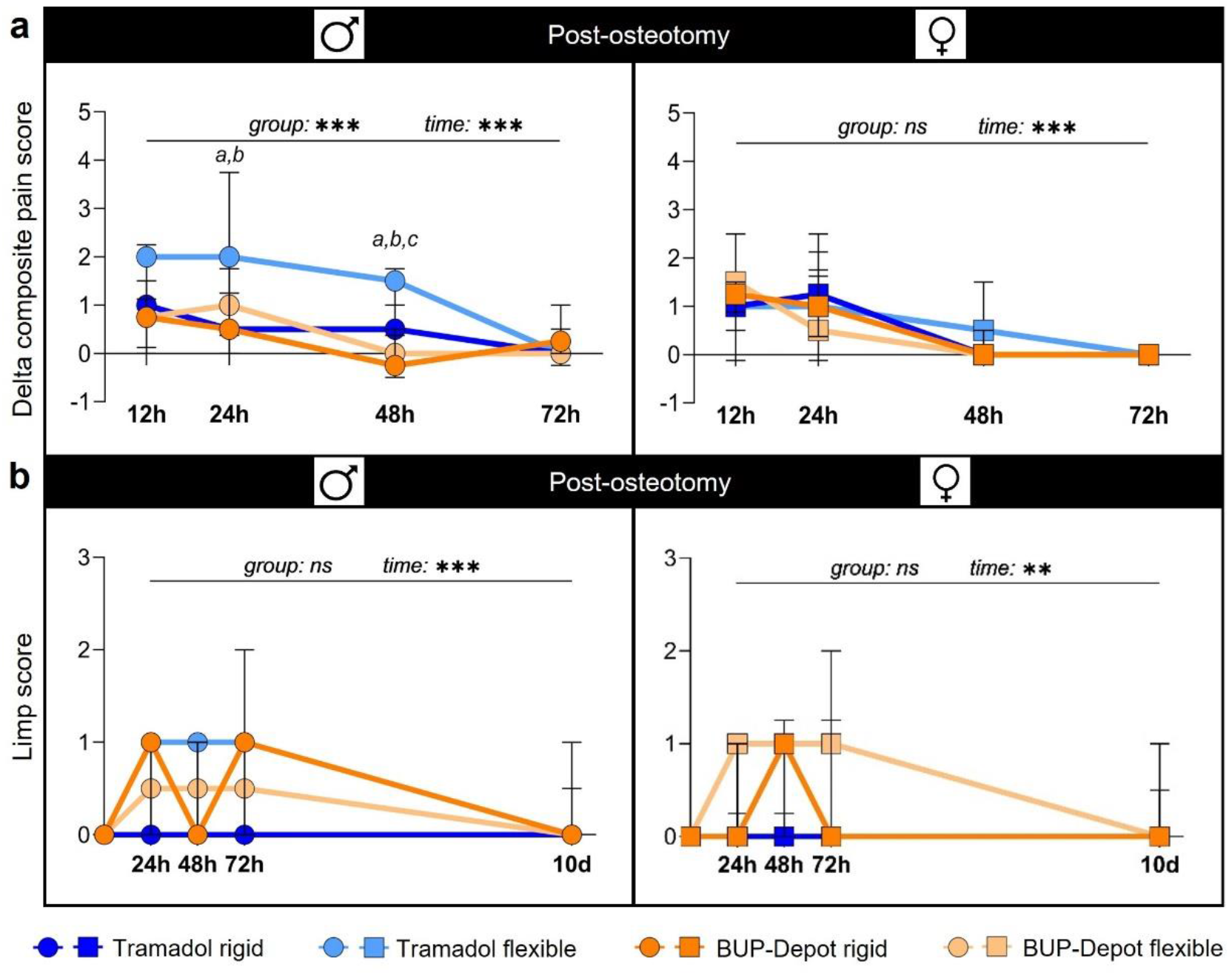
Delta composite pain score suggests adequate pain alleviation in most groups while limp score shows slight differences between groups post-osteotomy. (**a**) The delta composite pain score was evaluated at 12h, 24h, 48h and 72h post-anesthesia and post-osteotomy. For the delta composite score, scores from each individual mouse post-anesthesia were subtracted from their respective scores post-osteotomy. (**b**) Limp score was assessed at 24h, 48h, 72h and 10d post-osteotomy. All graphs show median with interquartile range for n = 8–10 (delta composite pain score) and n = 9–10 (limp score). Non-parametric ANOVA-type test - main effects of time and of group are represented in the graphs; exact *p-values* are listed in Table S5-S5.3 and Table S6; *\*p < 0.05, ***p < 0.001*. To determine group differences Kruskal-Wallis test and Dunn’s posthoc test with Bonferroni correction were performed. *a* – significant difference Tramadol rigid vs. Tramadol flexible; *b* – significant difference Tramadol flexible vs. BUP-Depot rigid; c – significant difference Tramadol flexible vs. BUP-Depot flexible.

### Limping indicates model-related alterations in walking behavior with only slight differences between groups

Walking behavior was assessed by i) using a metric limping score applied to individual 3 min videos and ii) analyses of gait and locomotion using the Noldus CatWalk XT at 24h, 48h, 72h, and 10d post-osteotomy. Walking behavior was also videotaped at the respective time points post anesthesia, but limping was only considered post-osteotomy, as mice did not show any alterations in walking post-anesthesia during routine monitoring and videotaping.

As expected, limping was observed at all time points post-osteotomy until 10d with a general improving trend (significant main time effect in males *p<0.001* and females *p= 0.018;* **Fig. 4b**; **Table S6**). In general, median scores ranged between 0 – 1 at 24h and 48h independent of sex and analgesics, while higher variations were seen at 72h (male BUP-Depot rigid; female BUP-Depot flexible; **Fig. 4b**). Slight alterations in limping were still visible after 10d in some individual mice in the female BUP-Depot groups and male Tramadol groups (all = score 1; sporadic limping; up to two animals, respectively).

### Gait and locomotion analysis indicates alterations in walking behavior and velocity with differences between groups

Analyses of gait and locomotion by the CatWalk XT exhibited a clear reduction in velocity after osteotomy in all groups (significant main time effect post-osteotomy for both sexes *p<0.001;* **Fig. 5a, b**; **Table S7**) with an upward trend towards 10d. At 24 and 48h post osteotomy, males with flexible fixation and tramadol treatment showed a significantly lower relative velocity compared to males with rigid fixation and BUP-Depot (non-parametric ANOVA-type test for group *p= 0.034;* Kruskal-Wallis test 24h *p= 0.052* and 48h *p= 0.038;* Dunn’s posthoc test – *p<0.05* when compared to BUP-Depot rigid; **Fig. 5a**; **Table S7.1–S7.3**). In females, significant differences in velocity were also evident at 24h post osteotomy, as mice with rigidly stabilized osteotomies exhibited elevated velocity compared to all other groups (non-parametric ANOVA-type test for group *p= 0.035;* Kruskal-Wallis test 24h *p= 0.009;* **Fig. 5b**; **Table S7.4–S7.5**). With respect to the relative mean intensity (a measure for load bearing) and relative stand duration, we did not observe significant group effects independent of sex and analgesic (**Fig. 5c–f**), while time-dependent reductions at 24h, 48h and 72h improved over time (relative mean intensity: non-parametric ANOVA-type test *p*< 0.001 in both sexes; relative stand duration *p< 0.001* in males and *p= 0.005* in females; **Table S8–S9**). However, the male mice with flexible fixation and Tramadol treatment showed numerically reduced values in relative mean intensity (**Fig. 5c**) when compared to the other males. In female mice, the relative stand duration only slightly increased over 10 days (median range 10d: 0.68 – 0.75; **Fig. 5f)**. The stride length was only markedly modulated over the first 72h but then recovered to baseline values at 10d (**Fig. 5g, h**; **Table S10**). Male mice with flexible fixation and Tramadol treatment also showed shortened stride length which was significant at 24h compared to BUP-Depot rigid, but not to Tramadol rigid or BUP-Depot flexible (non-parametric ANOVA-type test *p= 0.024;* Kruskal-Wallis test 24h *p= 0.044*; Dunn’s posthoc test – *p<0.05* when compared to BUP-Depot rigid; **Fig. 5g**; **Table S10-S10.2**), and did not recover over 10d. To evaluate the influence of the velocity on mean intensity, stand duration and stride length, we performed Spearman correlation analyses (**Fig. S5a–c**) which indicated that relative mean intensity (males r= 0.602, females r= 0.336; **Fig. S5a**) and relative stride length (males r= 0.801, females r= 0.522; **Fig. S5c**) correlated with the relative velocity (all *p< 0.001*) after osteotomy. Spearman correlation analyses showed only a very week correlation between relative stand duration and relative velocity (males r= −0.15, females r= 0.046) (both *p< 0.001*). Interpretation of those parameters must therefore be considered in context of the overall velocity.

**Figure 5.**
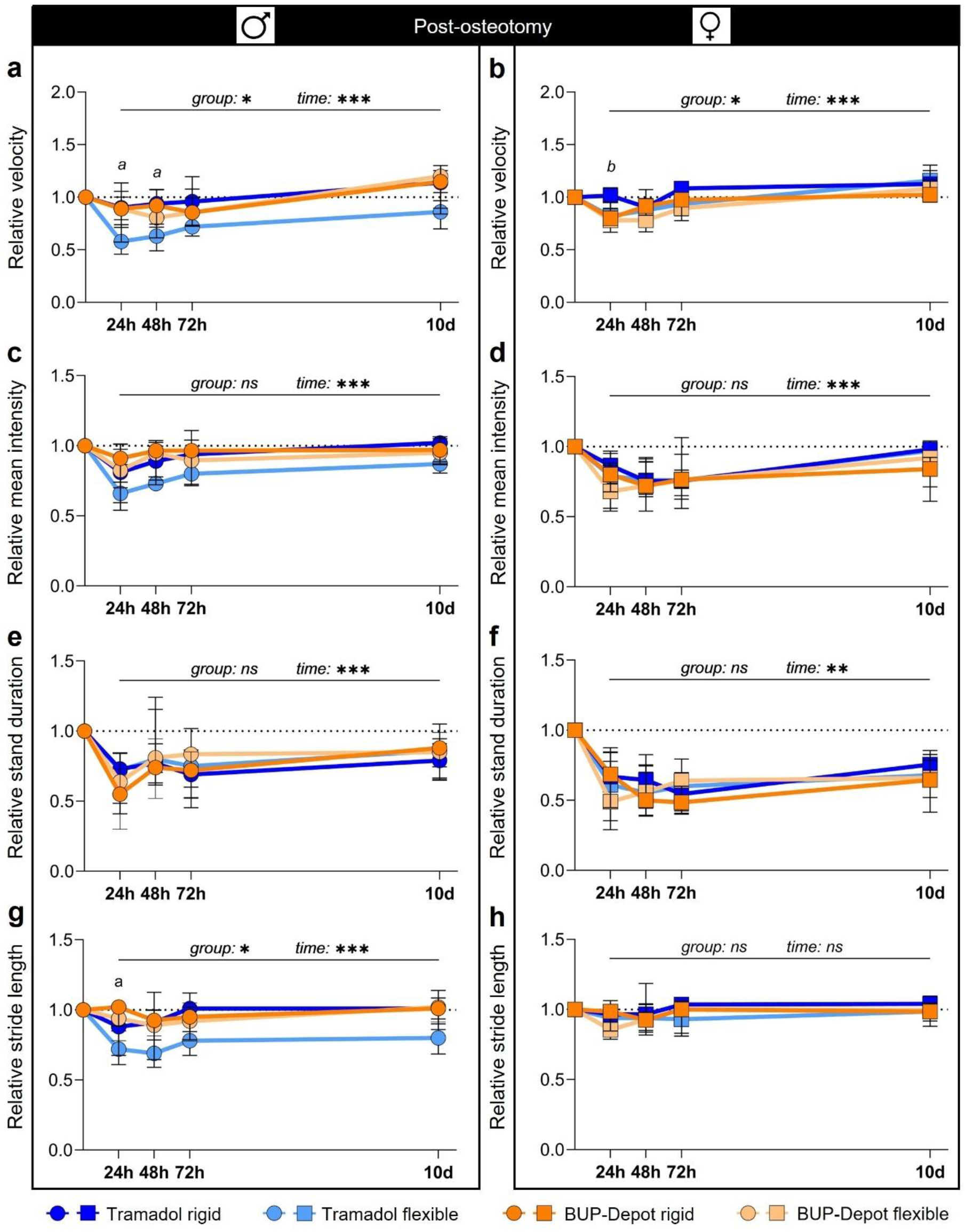
Gait analyses show model-related alterations in walking behavior specifically pronounced in male mice with flexible fixation and Tramadol treatment. CatWalk analysis was conducted at 24h, 48h, 72h and 10d post-osteotomy and normalized to the initial mean baseline value focusing on (**a, b**) relative velocity, (**c, d**) relative mean intensity, (**e, f**) relative stand duration and (**g, h**) relative stride length. All graphs show median with interquartile range for n = 8–10. Non-parametric ANOVA-type test - main effects of time and of group are represented in the graphs; exact *p-values* are listed in Table S7–S10.2; *\*p < 0.05, ***p < 0.001*. To determine group differences Kruskal-Wallis test and Dunn’s posthoc test with Bonferroni correction were performed. *a* – significant difference Tramadol flexible vs. BUP-Depot rigid; *b* – significant difference Tramadol rigid vs. BUP-Depot flexible.

### Fracture healing outcome is affected by fixation stability but not by analgesic regime

To evaluate potential effects of the BUP-Depot on fracture healing outcomes, we performed *ex-vivo* μCT (3-D), histomorphometric analysis (2-D) and vessel staining at day 14 post-osteotomy (**Fig. 6**). We observed a numerically lower BV/TV when comparing flexible fixation to rigid, except for males treated with Tramadol. However, the observed numerical differences did not reach statistical significance (BV/TV; **Fig. 6a, B**; **Fig. S6**; **Table S11**). The differences in BV/TV were slightly more pronounced in the BUP-Depot groups (male: median rigid = 25.87% vs. median flexible = 17.80%; female: median rigid = 26.12% vs. median flexible = 20.58%) than in the Tramadol groups (male: median rigid = 21.89% vs. median flexible = 21.52%; female: median rigid = 30.32% vs. median flexible = 23.31%).

**Figure 6.**
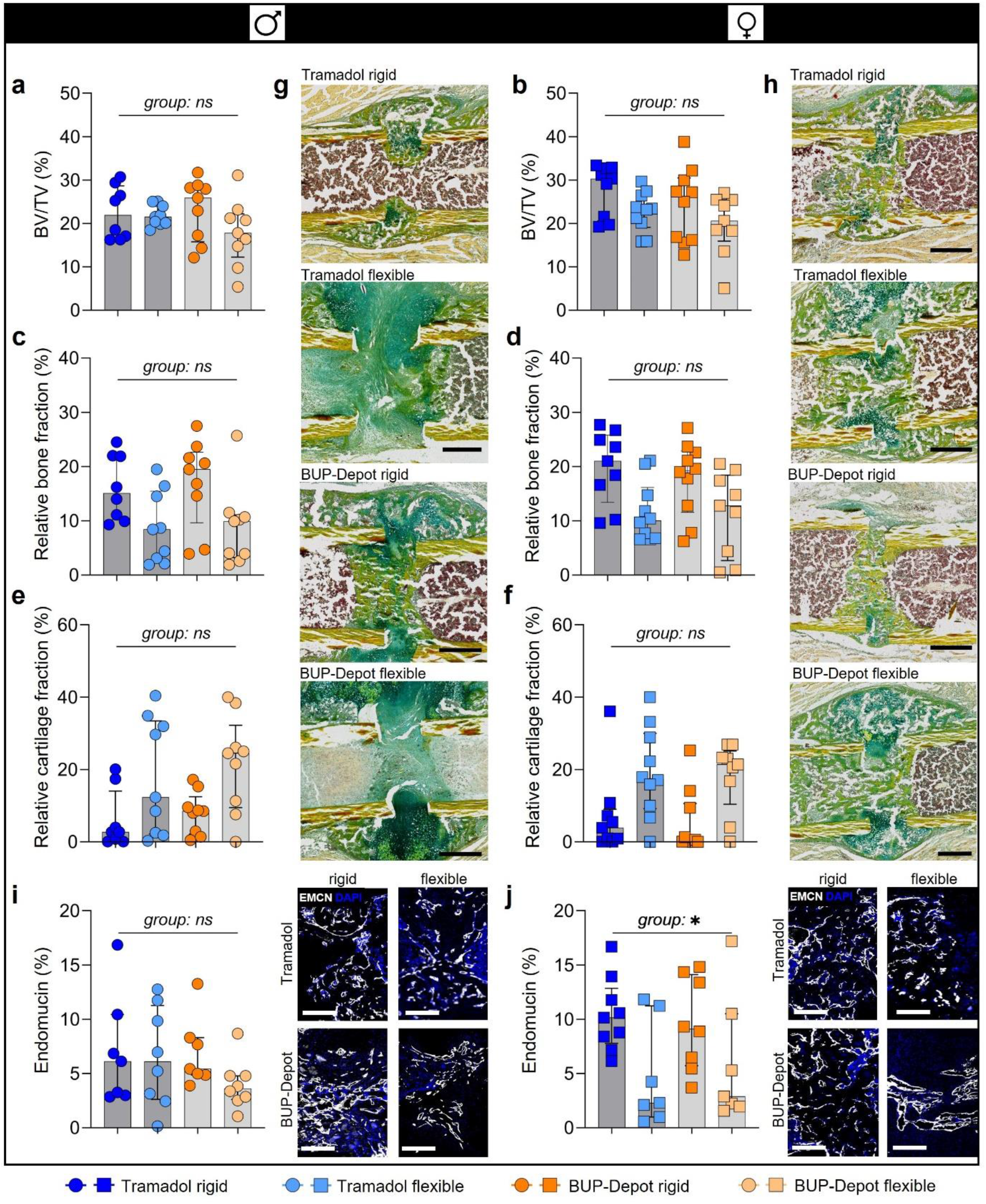
Analgesic regimes did not negatively affect fracture healing outcome at day 14, while the different fixations lead to differences in new bone formation. (**a, b**) Relative bone volume (BV/TV) (%), (**c, d**) relative bone fraction (%) and (**e, f**) relative cartilage fraction (%). (**g, h**) Exemplary images of the Movat’s pentachrome staining: yellow = mineralized bone, green = cartilage, magenta = bone marrow; scale bar 500 μm. (**i, j**) Immunofluorescence staining of vessel formation (Endomucin) including quantification and exemplary images; scale bar 200 μm. All graphs show median with interquartile range for n = 8–10 (**a-h**) and n = 6–9 (**i-j**). To determine group differences, Kruskal-Wallis test and Dunn’s posthoc test with Bonferroni correction were performed; exact *p-values* are listed in Table S11–S13.1; *\*p < 0.05*.

Histomorphometric analysis revealed comparable differences between the rigid and flexible fixation in relative bone and cartilage fraction. As expected, bone formation was reduced while cartilage formation was elevated in the flexible groups compared to rigid fixation in both sexes and treatment groups (**Fig 6c–h**; **Table S12**). Analysis of vessel formation (Endomucin/Emcn staining) within the callus area did not show significant or obvious differences in the male groups (**Fig. 6i**; **Fig. S6**; **Table S13**). Differences in vessel formation (Emcn) were shown between fixation in female mice (Kruskal-Wallis test *p= 0.045*; Dunn’s posthoc test *p> 0.05* between all groups; **Fig. 6j**; **Table S13.1**) with flexible fixation groups showing less relative Emcn^+^ areas when compared to the rigid fixation, independent of analgesia. Furthermore, DAPI staining indicated lower cellularity in the female mice with flexible fixation compared to rigid, also independent of analgesic treatment (**Fig. S6**).

## Discussion

In the present study, we evaluated the analgesic efficacy and possible side-effects of a newly developed sustained-release Buprenorphine (BUP-Depot) in comparison to an already established protocol, Tramadol in the drinking water, in two mouse-osteotomy models with different fixation stiffnesses ^19,23,24,26^. Due to individual differences in behavioral changes and pain perception ^19,27^, we chose a consecutive study design to analyze the effect of anesthesia and analgesia alone and in combination with an osteotomy in the same animal. The BUP-Depot delivered reliable pain relief over 72h post-surgical without side effects on fracture healing outcome, comparable to Tramadol applied with the drinking water.

General clinical parameters such as body weight, and food and water intake were noticeably reduced post-anesthesia and post-osteotomy. Reduction of food intake and therefore negatively influenced body weight development are known side-effects post-surgical but can also be related to anesthesia or Buprenorphine/Tramadol administration ^15,28–30^. Reduction of body weight observed after osteotomy was quite low (in the range of 5%) and similar to the anesthesia only control, which is indicative of an excellent pain management and wellbeing in all animals. Lowest values in body weight and food intake were reached after 24h in all groups, with an increase over 48h and 72h post-anesthesia and post-osteotomy. Interestingly, body weight loss at 24h was less pronounced after osteotomy than after anesthesia. The mice were 10 weeks of age during the first intervention and 12 weeks old at osteotomy. Mice show a rapid body weight development until skeletal maturity between approximately 10-12 weeks, which also includes the formation of more body fat ^31,32^. Since interventions entailing anesthesia result in short-term starvation during the recovery period, it can be speculated that mice at 10 weeks lost body weight more rapidly due to limited body fat reserves when compared to more mature mice (12 weeks). In general, the BUP-Depot showed similar effects on the body weight development as well as food and water intake as the established Tramadol treatment. We did not find differences in water intake between groups, excluding a negative effect of the Tramadol containing water on the overall drinking amount and ensuring a continuous uptake of medication as shown previously ^19,33^. However, the measurement of the overall 24h water intake does not allow to verify sufficient water intake during the first hours after surgery which might still be reduced as also reported by Evangelista et al. ^16^. To improve the food intake, food can be alternatively provided on the cage floor to prevent the animals from having to stand on their hind legs ^34^. However, this was not possible in this study. The use of high-caloric dietary gels could also be a valid alternative ensuring the consumption of food and liquids and can be also used as route for oral analgesic administration ^35^.

Changes in nest building and the willingness to explore foreign objects can indicate alterations in well-being in laboratory mice ^19,36–39^. In this study, nest building behavior was only scarcely influenced by anesthesia and osteotomy in all groups, independent of the analgesic regime, sex or fixation stiffness. This is in line with other studies suggesting that nest complexity scoring might not be sensitive enough to differentiate between minor pain and other potential stressors or reduced wellbeing after surgery ^37^. However, these findings are contrary to our previous study where we have reported a reduction in the nest building performance after osteotomy ^19^. Technical variances (amount of nesting material) or individual differences in scoring might explain the variations in our findings and underline the necessity for more objective approaches. The explorative behavior in male mice seemed to be negatively impacted by osteotomy, especially at 24h post-surgery, when compared to the post-anesthesia and female mice. Hohlbaum et al. also showed that female C57BL/6JRj mice exhibited shorter latency to explore than male mice 1 day after the last anesthesia in a repeated inhalation anesthesia trial, indicating a sex-specific difference that is in accordance with our findings with respect to explorative behavior ^40^.

A composite pain score was used combining the assessment of facial expression (parts of the mouse grimace scale) ^41^, and clinical appearance post-anesthesia and post-osteotomy. Based on our consecutive study design, we were able to calculate a delta for each individual mouse representing numerically the actual osteotomy effect. In line with other studies, the composite pain score was already slightly impacted by anesthesia and analgesia alone, indicating that some components of the composite pain score might not only be influenced by pain but rather also depict stress or discomfort ^19,42,43^. The highest delta composite scores were reached 12h and 24h after osteotomy suggesting the pain/discomfort peak due to the surgical procedure. However, median delta scores varied around 1, indicating only limited residual pain and/or discomfort over the first 48h which constantly declined till 72h post-surgical. The BUP-Depot provided comparable and sufficient alleviation of pain signs to the Tramadol treatment and our data indicate that pain relief can even be achieved over 72h after a single BUP-Depot injection. This is an important advantage compared to the commercially available sustained-release formulation Buprenorphine SR-LAB (ZooPharm – USA only), which yields effective serum concentrations over a maximum of 24h to 48h ^44,45^ with a potential need for re-administration if clinical signs indicate insufficient pain amelioration.

As model-specific parameters, we assessed limping behavior and locomotion using gait analysis. After osteotomy, limping was observed over 72h post-osteotomy in all groups, irrespective of sex, fixation, and analgesic, with no significant differences between the groups. More detailed gait analysis using the CatWalk system revealed reduced velocity and altered gait patterns over 72h and up to 10 days. While velocity, mean intensity and stride length ameliorated over 10 days in almost all groups (except male mice with flexible fixation and Tramadol analgesia), stand duration remained reduced in all groups. When analyzing CatWalk data, it needs to be considered that most gait parameters correlate to velocity and failing to address possible changes in velocity can affect the outcome of gait related data ^46–48^. In this study for example, relative velocity in male mice correlated strongly with relative mean intensity and stride length, but only weakly with other parameters which explains the comparable improvements over time. However, as the stand duration seems to be independent of the velocity, a limited functionality especially with regard to the full restoration of the musculature might be plausible explanation. As the type of surgery performed here requires splitting of the muscle and transection of the muscular insertion at the trochanter major, a certain degree of the observed gait alterations might be due to the not yet fully restored muscular function. In addition, limited mobility and changed gait are also common in human patients with e.g., proximal femur fractures of the femoral diaphysis and are not directly related to pain ^50,51^. Thus, we propose that the gait alteration in terms of stand duration over 10 days was likely caused by an unfinished functional restoration rather than pain or discomfort which is also supported by the absence of any additional pain-indicative signs at 72h. However, as the relative velocity was markedly reduced in male mice with the more flexible fixation and Tramadol as their analgesics, the reduced values in the relative mean intensity and the relative stride length in this group are most likely explained by the reduced velocity. This group also displayed the highest delta composite pain scores after 24 and 48h, although still in a low range. Because of the rather low pain scores it can be assumed, that peri- and post-surgical analgesia was sufficient, but could be improved, as there was a clinically relevant level of discomfort or pain. An explanation could be that the effective Tramadol dose was not achieved by application of 0.1 mg/g Tramadol due to the higher body weight of male mice. Evangelista et al. showed that male mice had lower serum concentrations of Tramadol than female mice when applying Tramadol (0.2 mg/ml) in the drinking water for up to 30h, although the analgesic effective M1 metabolite was similar between male and female mice ^33^. This is in line with a previous study performed in rats ^52^. In contrast, other studies report lower sensitivity of female mice or rats to Tramadol ^53,54^. With respect to the influence of the body weight on the local degree of interfragmentary movement, which might cause discomfort when too high, Röntgen et al. characterized two configurations of the external fixator, the rigid one (18.1 N/mm) and a very flexible one (0.82 N/mm), calculating the interfragmentary strain in a 0.5 mm osteotomy gap for a 25 g mouse with 2.8% and 61%, respectively ^55^. However, when applied to female C57BL/6 mice, they did not find differences in body weight, ground reaction force and locomotion over 18 days, indicating sufficient analgesia even in the presence of higher local strains ^55^. These observations highlight potential sex-specific differences in pain perception but also the response to analgesic medication ^56^. Sex-specific adaptions of pain management regimes are therefore advisable in future studies. Our findings underline further, that pain management in animals experiments requires a constant reevaluation of the chosen protocol as well as the consideration of strain, sex, interindividual differences in animals, procedure, options to reduce re-injections and more ^6^. As not only sex, but also genetics influences experienced pain and response to analgesia in mice ^56,57^, further studies using different strains are needed in the evaluation of commonly used analgesic regimens in laboratory rodents.

In terms of fracture healing outcome analyzed by *ex-vivo* μCT and histomorphometry, we found no differences between the analgesic regimens, indicating safe use of the newly developed BUP-Depot in the analyzed models. A more flexible fixation allows higher interfragmentary movements and, therefore, promotes cartilage rather than bone formation as well as formation of a larger periosteal callus ^55^, as seen in the histomorphometric analysis. Staining for endomucin, representing vessels, revealed no difference between fixations or analgesic in male mice. However, female mice with more flexible stabilized osteotomies showed a reduced Emcn-positive area as well as lower cellularity, regardless of their analgesic regime. Besides mechanical hindrance of revascularization due to higher strains, the higher proportion of cartilage observed in female mice with flexible stabilized osteotomies might have prevented revascularization due to the intrinsic anti-angiogenic nature of cartilage e.g., due to chondromodulin-1 ^58,59^. This reduced vascularization is in accordance with earlier observations in sheep ^60^. The different patterns of Emcn staining in males and females might also hint on a more advanced callus remodeling and therefore a more rapid healing progression in males than females ^61^.

An effective sustained-release Buprenorphine in Europe would not only be a conceivable alternative to the application of Tramadol with the drinking water as investigated in this study, but also a potential alternative to repeated injections of Buprenorphine, which are still most frequently used for pain management in femur fracture models ^9^. Assessment of the analgesic effect of Buprenorphine is most often based measurement of plasma or blood serum levels. Studies specified therapeutic effective concentrations of Buprenorphine in plasma at a threshold of around 1 ng/ml or 1 ng/g in mice and rats ^10,65,66^. However, Buprenorphine works through the μ-, κ- and δ-opioid receptors in the brain ^67,68^ and it can be hypothesized that reliable pain alleviation is more reliant on specific binding concentration values in the brain than specific plasma concentrations ^23^, as demonstrated by a correlation between analgesic effects and specific binding concentrations of Buprenorphine in the brain of rats ^69^. Schreiner et al. contemplated that specific binding concentrations of 5 ng/g (as observed 24h after injection of the BUP-Depot) in the brain might be needed for reliable pain relieve in mice. Binding concentrations of less than 3 ng/g at 48h post injection of BUP-Depot – a concentration comparable to levels observed 12h after Temgesic injection – still resulted in high, but not significantly increased withdrawal latencies compared to a single injection with Temgesic or NaCl ^23^. They therefore suggest that alleviation of strong pain through the BUP-Depot might require the administration every 24h ^23^. However, based on our assessment of the clinical, behavioral, and model-specific parameters, we can postulate that the analgesic properties of the BUP-Depot are sufficient for 72h post-operative analgesia in our specific mouse-osteotomy model of moderate severity. Nonetheless, the potential use and the possible need for re-application of the BUP-Depot in other, more painful models still needs to be critically assessed and evaluated.

Taken together, our assessment of clinical, behavioral, and model-specific parameters suggest that the analgesic properties of the BUP-Depot were sufficient for 72h post-operative analgesia in male and female C57BL/6N mice after femoral osteotomy stabilzed with external fixators. The BUP-Depot therefore provides an excellent alternative for extended pain relief in preclinical studies. The availability of such a sustained-release formulation of Buprenorphine in Europe would be substantially beneficial for mouse analgesia in animal experiments.

## Animals, Material and Methods

### Ethics and guidelines

The study was conducted according to the guidelines of the German Animal Welfare Act, National Animal Welfare Guidelines, and was approved by the local Berlin state authority (Landesamt für Gesundheit und Soziales – LAGeSo; permit number: G0044/20). Health monitoring in the animal facility was performed according to the FELASA guidelines (Supplementary Information).

### Animals and husbandry

A total of 40 male and 40 female C57BL/6N mice aged 8 weeks were either provided by the Experimental Medicine Research Facilities (Charité - Universitätsmedizin Berlin, Berlin, Germany) or purchased from Charles River Laboratories (Sulzfeld, Germany). Mice underwent the first intervention (anesthesia/analgesia) with 10 weeks (body weight – males: 25.96 ± 2.1 g; females: 21.41 ± 1.3 g) and osteotomy with 12 weeks (body weight – males: 27.61 ± 1.9 g; females: 22.75 ± 1.3 g). Mice were housed in a semi-barrier facility in individually ventilated cages (IVC, Eurostandard Type II, Tecniplast, Milan, Italy). Housing conditions encompassed a 12/12–h light/dark cycle (light from 6:00 a.m. to 6:00 p.m.), room temperature of 22±2°C and a humidity of 55±10%. Food (Standard mouse diet, Ssniff Spezialdiäten, Soest, Germany) and tap water were available *ad libitum*.

Mice were randomly divided into groups of two per cage. If mice had to be separated due to aggressive behavior, they were housed in a separated pair housing system in Green Line IVC Sealsafe PLUS Rat GR 900 cages (Tecniplast, Milan, Italy), which were divided in two equally sized compartments by a perforated transparent partition wall. Cages contained wooden chips (SAFE FS 14, Safe Bedding, Rosenberg, Germany), 20 g Envirodri (Shepherd Specialty Papers, USA), and a shredded paper towel as bedding and nesting material, a clear handling tube (Datesand Group, Bredbury, UK) and a mouse double swing (Datesand Group, Bredbury, UK). No houses were provided to allow reliable scoring and to reduce the risk of injury after osteotomy. After osteotomy, tunnels and double swings were removed from the cages to reduce the risk of injury. Two single swings (Datesand Group, Bredbury, UK) per cage were reinstalled 5 days after osteotomy. Animals were tunnel handled only and all experimenters performing analyses were female. Animal husbandry and care were in accordance with contemporary best practice.

### Study design and experimental timeline

Reporting of this study was carried out in compliance with the ARRIVE 2.0 guidelines, including the “Arrive Essential 10” and most of the “Arrive Recommended Set”. The study was pre-registered in the Animal Study Registry (Bf3R, Germany; DOI: 10.17590/asr.0000221).

The study included 8 groups with each n= 9–10 mice, comparing male and female mice, rigid and flexible external fixators, and two different pain management protocols: tramadol in drinking water or sustained-release Buprenorphine (BUP-Depot) (**Fig. 1a**). Cages and mice received individual random numbers that did not allow any inferences for the analgesic regimen or fixation group. Experimenters performing the pre- and post-surgical training and investigations were blinded.

After acclimation for 5 days, training was performed by one female experimenter, following a 2-week schedule to accustom the mice to the experimenter, tunnel handling, Noldus CatWalk XT (Noldus, Wageningen, Netherlands) and observation boxes (Ugo Basile, Gemonio, Italy). Baseline measurements were also obtained during this period (**Fig. 1b**). To correct for individual behavioral changes induced by anesthesia and analgesia alone, mice were first anesthetized and received their assigned analgesic protocol without any further surgical procedure (first intervention). Then, parameters were assessed at 12h, 24h, 48h, and 72h post-procedure. 14 days after the first intervention, the same animals were subjected to anesthesia, analgesia, and osteotomy (second intervention) and assessed at the above listed time points as well as at day 10 post-surgical. Mice were euthanized 14 days post-osteotomy to retrieve the osteotomized femur.

### Analgesic regimes

Each mouse received one s.c. injection of regular Buprenorphine (1 mg/kg Temgesic, RB Pharmaceuticals, Heidelberg, Germany) at the beginning of each intervention (anesthesia/analgesia alone and osteotomy). Depending on the randomly assigned group, mice either additionally received Tramadol administered with the drinking water (0.1 mg/ml, Tramal Drops, Grünenthal, Stolberg, Germany) or a s.c. injection of the BUP-Depot (1.2 mg/kg). Tramadol was administered one day before and three consecutive days after both interventions. The BUP-Depot was injected once at the end of both interventions. BUP-Depot (RG 502 H-Big) was prepared at the University of Basel, Switzerland, as described previously by Schreiner et al. ^23,24^. Four different batches of BUP-Depot were imported in accordance with national regulations for controlled substances (BtM import authorization No. 4679477). Each batch was analyzed prior to shipment for drug content, reconstitution time, and drug release kinetics as described previously ^23,24^. The BUP-Depot was stored as a lyophilizate in glass vials at 4°C. Each vial was reconstituted with physiological saline (0.9% NaCl) immediately before administration.

### Anesthesia and osteotomy

Independent of the intervention, all mice were anesthetized with isoflurane (~2–3%; provided in 100% oxygen; CP-Pharma, Burgdorf, Germany) before being weighed and moved onto a heating pad (37°C). Anesthesia was maintained at ~2% via a nose cone. Eye ointment, physiological saline (0.5 ml, 0.9% NaCl), Clindamycin (45 mg/kg, Ratiopharm, Ulm, Germany) and a single s.c. injection of Buprenorphine were applied. Anesthesia was then upheld for 15 minutes for the first intervention (anesthesia and analgesia alone). For osteotomy, the left femur was shaved and disinfected with alcoholic iodine solution. The osteotomy was conducted as described previously ^19,70,71^. A longitudinal skin incision was made between knee and hip, and the musculus vastus lateralis and musculus biceps femoris were bluntly separated to expose the femur. Two standardized external fixators (rigid: 18.1 N/mm; flexible: 3.2 N/mm, both RISystem, Davos, Switzerland) were used for stabilization. The external bar of the fixator was positioned parallel to the femur and all pins were positioned accordingly. Afterwards, an approximately 0.5 mm osteotomy gap was created between the second and third pin using a Gigli wire saw (0.44 mm; RISystem, Davos, Switzerland) and the gap was flushed with saline. Muscle and skin were closed with two layers of sutures (muscle: 5-0 Vicryl, skin: Ethilon 5-0, both Ethicon, Raritan, USA). For recovery, the mice were returned to their home cages under an infrared lamp and were closely monitored.

### Body weight and food/water intake

Animals were weighed before the intervention (defined as 0h), and at 12h, 24h, 48h and 72h after both interventions and then every other day until osteotomy/euthanasia. Food/water intake were measured per cage (i.e., two mice) by weighing the food and water bottles every 24h, beginning 1 day prior to each intervention and ending 3 days after. The difference to the previous value was calculated. All measurements were normalized to the respective baseline values at time point 0h.

### Explorative test and nest complexity score

Both scores were assessed before any other assessment or handling of the mice. To examine the motivation of the mice to explore and interact (sniffing, holding with forepaws, or carrying) with a foreign object, we added a Nestlet (Ancare, Bellmore, USA) to the home cages and observed the mice for one minute. The explorative test was scored 1 (interaction) or 0 (no interaction) per cage. An interaction of one animal of the cage was deemed as sufficient for a positive score. Nest complexity scoring was performed following Hess et al. ^72^ in the home cage assigning scores between 0 – 5.

### Composite pain score

A composite pain score was used to combine the assessment of facial expression and clinical appearance ^19,41^ (**Table 1**). The maximal score was 9. At 12h, 24h, 48h, and 72h post-intervention, the mice were transferred into a clear observation box and individually filmed for 3 min after an acclimatization period of 1 min (Basler Video Recording Software, Ahrensburg, Germany). Video analysis was performed by one blinded observer. As anesthesia and analgesia alone also affect the composite pain score, we calculated the delta composite pain score for each mouse by subtracting the scores from each individual mouse post-anesthesia from their respective scores post-osteotomy. This allowed us to evaluate the effect of the surgical procedure on an individual base without the interference of behavioral or clinical changes induced by the anesthesia and analgesia alone ^73^.

**Table 1:** Composite pain score.

| Parameter | Specification | Score<br>(0 = not present) |
| --- | --- | --- |
| Orbital tightening | Faint narrowing of the orbital area up to a tightly closed eyelid | Range: 0 – 2<br>0.5 = minimal<br>1 = moderately<br>1.5 = moderately severe<br>2 = severe |
| Ear position | Ears pulled back or rotated outwards and/or backwards | Range: 0 – 2<br>1 = moderately<br>2 = severe |
| Posture | Crouched posture, head and nose positioned towards the ground | Range: 0 – 1<br>0.5 = held for less than 10 s<br>1 = held for more than 10 s |
| Coat condition | Coat appeared disheveled or unkempt | Range: 0 – 1<br>0.5 = only certain body parts<br>1 = all over the body |
|  | Erected fur, mice appear scruffy | Range: 0 – 1<br>0.5 = only on one body part<br>1 = on more than one body part, generalized |
| Movement | Apathetic, sedated, tipsy (0 – 1);<br>Crawling (0 – 1) | Range: 0 – 1 each<br>0.5 = minimal<br>1 = moderately |

### Walking behavior – limp score

To assess the walking behavior of each mouse, limp scoring was performed as described by Jirkof et al. ^19^. The mice were transferred to conventional type III cages that contained the same type of wooden chips as the home cages. After an acclimation period, a 3 min video was recorded. Walking behavior was examined at time points concurrent with the CatWalk analysis at 24h, 48h, and 72h post-anesthesia and post-osteotomy as well as 10d post-osteotomy. Video analysis was performed by two blinded observers and scores from 0 – 4 were assigned (**Table 2**). If walking seemed to be impaired due to a mechanical problem (e.g., displacement of the patella) one point was subtracted from the assigned score.

**Table 2:** Limp score.

| Specification | Limp Score |
| --- | --- |
| Normal use, physiological gait | 0 |
| Complete ground contact and sporadic limping or alteration of the gait pattern | 1 |
| No complete ground contact or limping, constant alteration of the gait pattern | 2 |
| Partial non-use of the limb | 3 |
| Complete lack of use | 4 |

### CatWalk analysis

Specific gait analysis was performed using the CatWalk XT Gait Analysis system for rodents. Multiple runs per animal were acquired before interventions and at 24h, 48h, 72h post-anesthesia (data not shown in results) and post-osteotomy, as well as 10d post-osteotomy. Post-acquisition the runs were screened, and non-compliable runs as well as interrupted runs (e.g., by sniffing, rearing) were excluded. Runs that noticeably differed from the rest of the runs of this trial were also excluded, leaving an average of 4.4 runs per animal and time point for analyses. All runs were classified automatically by the Noldus CatWalk XT software (version XT10.6) and revised for classification errors (i.e., incorrect identification of paws), which were corrected manually. From the obtained data, mean speed (cm/s) (velocity) and the following parameters were analyzed for the osteotomized left hind leg: mean intensity, stand duration (s) and stride length (cm). The two baseline measurements were used to calculate one baseline average value. Values at all time points were then normalized to the respective average baseline (time point measurement divided by average baseline value).

### Euthanasia and sample collection

Euthanasia was carried out according to contemporary best practice. At 14 days post-osteotomy, mice were euthanized by cervical dislocation in deep anesthesia. The osteotomized femora were retrieved and fixed in 4% paraformaldehyde (PFA; Electron Microscopy Sciences, Hatfield, USA) at 4°C for 6–8h. The femora were then transferred into PBS until ex-vivo μCT was completed.

### Ex-vivo μCT

To determine bone formation three-dimensionally, femurs were scanned in a SkyScan 1172 high-resolution μCT (Bruker, Kontich, Belgium). Voxel size was set to 8 μm and the bones were scanned with a source energy of 70 kV, 142 μA, a rotation step of 0.2 degree and an 0.5 mm aluminum filter. Scans were reconstructed using NRecon (Bruker, Kontich, Belgium), applying ring artefact reduction and beam hardening corrections. CT Analyser software (version 1.20.3.0; both Bruker, Kontich, Belgium) was used for 2D and 3D analyses. By excluding the original cortical bone within the callus, the total volume (TV, mm^3^), the total bone volume (BV, mm^3^) and the bone volume fraction (BV/TV) of the newly formed bone were analyzed in a manually defined volume of interest (VOI) ^18^.

### Histology and immunofluorescence

Following *ex-vivo* μCT, bones were placed in ascending sugar solutions as cryoprotectant (10%, 20%, 30%) at 4°C for 24h each, then cryo-embedded in SCEM medium (Sectionlab, Japan) and stored at - 80°C. Consecutive sections of 7 μm were prepared using a cryotome (Leica, Wetzlar, Germany) and cryotape (Cryofilm 2C(9), Sectionlab, Japan). Sections were fixed onto glass slides, air-dried, and stored at −80 °C until staining. Movat’s pentachrome staining comprised the following steps: sections were air dried for 15 min, fixed with 4% PFA (30 min; Electron Microscopy Sciences, Hatfield, USA), pretreated with 3% acetic acid for 3 min, stained 30 min in 1% alcian blue pH 2.5, followed by washing in 3% acetic acid under light microscopic control. Sections were rinsed in H_2_O_dest_ and immersed in alkaline ethanol for 60 min, then washed in tap water followed by incubation in Weigert’s hematoxylin for 15 min. After washing in tap water for 10 min, sections were stained in crocein scarlet-acid fuchsin for 15 min, treated with 0.5% acetic acid for 1 min, followed by 20 min incubation in 5% phosphotungstic acid, and 1 min in 0.5% acetic acid. The sections were washed three times for 2 min in 100% ethanol, followed by incubation in alcoholic Saffron du Gâtinais for 60 min. The slides were dehydrated in 100% ethanol, cleared shortly in xylene, covered with Vitro-Clud and a cover slip. Imaging was performed on a Leica light microscope using LAS X software (Leica Microsystems GmbH, Wetzlar, Germany) at 10x magnification. Quantitative analyses of the Movat’s pentachrome staining were evaluated using an ImageJ macro. All analyses were performed blinded to sex, fixation, and pain management protocol.

Immunofluorescence staining was performed as described previously ^70,71^ using the following antibody: Endomucin (Emcn) (V.7C7 unconjugated, rat monoclonal, sc-65495, 1:100; Santa Cruz Biotechnology, Dallas, USA), goat anti-rat A647 (1:500; A-21247, polyclonal, Invitrogen, Thermo Fisher Scientific, Waltham, USA) and DAPI (1:1,000; Thermo Fisher Scientific, Waltham, USA). Blocking was performed with 10% FCS/PBS and staining solution contained 5% FCS and 0.1% Tween20 (Sigma Aldrich, St. Louis, USA). Images were acquired using a Keyence BZ9000 microscope (Keyence, Osaka, Japan). The images were processed and analyzed with ImageJ ^74,75^. An area of interest was established and managed via the built-in ROI-Manager, while cell number and signal distribution within the area were determined using the plug-ins Cell-counter and Calculator Plus. Data was processed with the ImageJ plugin OriginPro.

### Statistical analysis

Sample size was calculated based on own preliminary data ^19^ using a nonparametric ranking procedure to analyze longitudinal data. The required number of animals was modeled in R using the package nparLD ^76^. Assuming a 20% difference and a power of ~80% resulted in n=10 animals per group.

Statistical analysis was performed using RStudio and graphs were created in GraphPad Prism (V9). To test whether the data from female and male mice are homogenous and would allow for an integrated data analysis, body weight data of both sexes was first representatively compared using the F2-LD-F1 design. Since we found significant interaction regarding sex, both sexes were analyzed separately for all statistical analyses. Nonparametric analysis of longitudinal data (F1-LD-F1 design; further on named non-parametric ANOVA-type tests), was used to test for significant differences in the main effect of time and main effect of group, separated by sex. When mean group differences *p≤0.05* were detected, group comparison for each time point was performed using the Kruskal-Wallis test ^77^. To determine group differences, Dunn’s posthoc test with Bonferroni correction was performed for each time point ^78,79^. Non-parametric ANOVA-type tests as well as exact *p-values*, chi-squared and df of all analyses are provided in the Supplementary Information. Excluded mice and data are detailed in the Supplementary Information.

## Conflict of interest statement

The authors declare that they have no conflict of interests.

## Author contributions

Study conception: A.L., A.R., P.J., J.H.; study conduction: A.W., A.L., A.R., V.S., C.B., K.H., S.K., R.K.; data collection: A.W., A.L., A.R., S.K.; analysis and interpretation: A.W., A.L., A.R., C.B., F.K.; drafting manuscript: A.W., A.L., A.R.; revising manuscript: C.B., K.H., M.L., C.T.-R., F.B., J.H., P.J.

## Competing interests

The authors declare no competing interests.

## Acknowledgement

We would like to thank Dr. Kristina Ullmann, Bianka Verrett and Sylvia Gundelach for their support during the planning and performance of the animal experiments. Special thanks to Darryl Borland and PD Dr. Maxim Puchkov for expert technical support and the preparation of BUP-Depot. This study was funded by Charite3R, Charité-Universitätsmedizin Berlin (RefineMOMo 2.0; RefineLab) and a grant from Interpharma Basel.

## Data availability statement

The authors declare that all data supporting the findings of this study are available within the paper and its Supplementary Information file. Further information is made available by the authors upon request.

## Notes

### Competing Interest Statement

The authors have declared no competing interest.

### Summary of Updates

manuscript restructured; discussion streamlined; Supplemental file updated

## References

1 Balcombe, J. P., Barnard, N. D. & Sandusky, C. Laboratory routines cause animal stress. Contemp Top Lab Anim Sci 43, 42–51 (2004).

2 Carstens, E. & Moberg, G. P. Recognizing pain and distress in laboratory animals. Ilar j 41, 62–71, doi:10.1093/ilar.41.2.62 (2000).

3 Guo, S. & Dipietro, L. A. Factors affecting wound healing. J Dent Res 89, 219–229, doi:10.1177/0022034509359125 (2010).

4 Page, G. G. The immune-suppressive effects of pain. Adv Exp Med Biol 521, 117–125, doi:https://www.ncbi.nlm.nih.gov/books/NBK6140/ (2003).

5 Peterson, N. C., Nunamaker, E. A. & Turner, P. V. To Treat or Not to Treat: The Effects of Pain on Experimental Parameters. Comp Med 67, 469–482 (2017).

6 Jirkof, P. Side effects of pain and analgesia in animal experimentation. Lab Anim (NY) 46, 123–128, doi:10.1038/laban.1216 (2017).

7 Flecknell, P. Rodent analgesia: Assessment and therapeutics. Vet J 232, 70–77, doi:10.1016/j.tvjl.2017.12.017 (2018).

8 Carbone, L. & Austin, J. Pain and Laboratory Animals: Publication Practices for Better Data Reproducibility and Better Animal Welfare. PLoS One 11, e0155001, doi:10.1371/journal.pone.0155001 (2016).

9 Wolter, A. et al. Systematic review on the reporting accuracy of experimental details in publications using mouse femoral fracture models. Bone 152, 116088, doi:10.1016/j.bone.2021.116088 (2021).

10 Guarnieri, M. et al. Safety and efficacy of buprenorphine for analgesia in laboratory mice and rats. Lab Anim (NY) 41, 337–343, doi:10.1038/laban.152 (2012).

11 Roughan, J. V. & Flecknell, P. A. Buprenorphine: a reappraisal of its antinociceptive effects and therapeutic use in alleviating post-operative pain in animals. Lab Anim 36, 322–343, doi:10.1258/002367702320162423 (2002).

12 Bullingham, R. E., McQuay, H. J., Moore, A. & Bennett, M. R. Buprenorphine kinetics. Clin Pharmacol Ther 28, 667–672, doi:10.1038/clpt.1980.219 (1980).

13 Gades, N. M., Danneman, P. J., Wixson, S. K. & Tolley, E. A. The magnitude and duration of the analgesic effect of morphine, butorphanol, and buprenorphine in rats and mice. Contemp Top Lab Anim Sci 39, 8–13 (2000).

14 Yu, S. et al. Pharmacokinetics of buprenorphine after intravenous administration in the mouse. J Am Assoc Lab Anim Sci 45, 12–16 (2006).

15 Jirkof, P., Tourvieille, A., Cinelli, P. & Arras, M. Buprenorphine for pain relief in mice: repeated injections vs sustained-release depot formulation. Lab Anim 49, 177–187, doi:10.1177/0023677214562849 (2015).

16 Evangelista-Vaz, R., Bergadano, A., Arras, M. & Jirkof, P. D. Analgesic Efficacy of Subcutaneous-Oral Dosage of Tramadol after Surgery in C57BL/6J Mice. J Am Assoc Lab Anim Sci 57, 368–375, doi:10.30802/AALAS-JAALAS-17-000118 (2018).

17 Girón, R. et al. Synthesis and opioid activity of new fentanyl analogs. Life Sci 71, 1023–1034, doi:10.1016/s0024-3205(02)01798-8 (2002).

18 Bucher, C. H. et al. Experience in the Adaptive Immunity Impacts Bone Homeostasis, Remodeling, and Healing. Front Immunol 10, 797, doi:10.3389/fimmu.2019.00797 (2019).

19 Jirkof, P. et al. Administration of Tramadol or Buprenorphine via the drinking water for post-operative analgesia in a mouse-osteotomy model. Sci Rep 9, 10749, doi:10.1038/s41598-019-47186-5 (2019).

20 Rapp, A. E. et al. Induced global deletion of glucocorticoid receptor impairs fracture healing. Faseb j 32, 2235–2245, doi:10.1096/fj.201700459RR (2018).

21 Histing, T. et al. Obesity does not affect the healing of femur fractures in mice. Injury 47, 1435–1444, doi:10.1016/j.injury.2016.04.030 (2016).

22 Sauer, M., Fleischmann, T., Lipiski, M., Arras, M. & Jirkof, P. Buprenorphine via drinking water and combined oral-injection protocols for pain relief in mice. Applied Animal Behaviour Science 185, 103–112, doi:10.1016/j.applanim.2016.09.009 (2016).

23 Schreiner, V. et al. Design and in vivo evaluation of a microparticulate depot formulation of buprenorphine for veterinary use. Sci Rep 10, 17295, doi:10.1038/s41598-020-74230-6 (2020).

24 Schreiner, V., Detampel, P., Jirkof, P., Puchkov, M. & Huwyler, J. Buprenorphine loaded PLGA microparticles: Characterization of a sustained-release formulation. Journal of Drug Delivery Science and Technology 63, doi:10.1016/j.jddst.2021.102558 (2021).

25 Webster, L. R., Camilleri, M. & Finn, A. Opioid-induced constipation: rationale for the role of norbuprenorphine in buprenorphine-treated individuals. Subst Abuse Rehabil 7, 81–86, doi:10.2147/SAR.S100998 (2016).

26 Rapp, A. E. et al. Analgesia via blockade of NGF/TrkA signaling does not influence fracture healing in mice. J Orthop Res 33, 1235–1241, doi:10.1002/jor.22892 (2015).

27 Hohlbaum, K. et al. Impact of repeated anesthesia with ketamine and xylazine on the well-being of C57BL/6JRj mice. PLoS One 13, e0203559, doi:10.1371/journal.pone.0203559 (2018).

28 Brennan, M. P., Sinusas, A. J., Horvath, T. L., Collins, J. G. & Harding, M. J. Correlation between body weight changes and postoperative pain in rats treated with meloxicam or buprenorphine. Lab Anim (NY) 38, 87–93, doi:10.1038/laban0309-87 (2009).

29 Clark, M. D. et al. Evaluation of liposome-encapsulated oxymorphone hydrochloride in mice after splenectomy. Comp Med 54, 558–563 (2004).

30 Goecke, J. C., Awad, H., Lawson, J. C. & Boivin, G. P. Evaluating postoperative analgesics in mice using telemetry. Comp Med 55, 37–44 (2005).

31 Ferguson, V. L., Ayers, R. A., Bateman, T. A. & Simske, S. J. Bone development and age-related bone loss in male C57BL/6J mice. Bone 33, 387–398, doi:https://doi.org/10.1016/S8756-3282(03)00199-6 (2003).

32 Yang, Y., Smith, D. L., Jr., Keating, K. D., Allison, D. B. & Nagy, T. R. Variations in body weight, food intake and body composition after long-term high-fat diet feeding in C57BL/6J mice. Obesity (Silver Spring, Md.) 22, 2147–2155, doi:10.1002/oby.20811 (2014).

33 Evangelista Vaz, R. et al. Preliminary pharmacokinetics of tramadol hydrochloride after administration via different routes in male and female B6 mice. Vet Anaesth Analg 45, 111–122, doi:10.1016/j.vaa.2016.09.007 (2018).

34 Lang, A., Schulz, A., Ellinghaus, A. & Schmidt-Bleek, K. Osteotomy models - the current status on pain scoring and management in small rodents. Lab Anim 50, 433–441, doi:10.1177/0023677216675007 (2016).

35 Nizar, J. M., Bouby, N., Bankir, L. & Bhalla, V. Improved protocols for the study of urinary electrolyte excretion and blood pressure in rodents: use of gel food and stepwise changes in diet composition. American Journal of Physiology-Renal Physiology 314, F1129–F1137, doi:10.1152/ajprenal.00474.2017 (2018).

36 Gaskill, B. N., Karas, A. Z., Garner, J. P. & Pritchett-Corning, K. R. Nest building as an indicator of health and welfare in laboratory mice. J Vis Exp, 51012, doi:10.3791/51012 (2013).

37 Jirkof, P. et al. Assessment of postsurgical distress and pain in laboratory mice by nest complexity scoring. Lab Anim 47, 153–161, doi:10.1177/0023677213475603 (2013).

38 Jirkof, P. Burrowing and nest building behavior as indicators of well-being in mice. J Neurosci Methods 234, 139–146, doi:10.1016/j.jneumeth.2014.02.001 (2014).

39 Rock, M. L. et al. The time-to-integrate-to-nest test as an indicator of wellbeing in laboratory mice. J Am Assoc Lab Anim Sci 53, 24–28 (2014).

40 Hohlbaum, K. et al. Severity classification of repeated isoflurane anesthesia in C57BL/6JRj mice-Assessing the degree of distress. PloS one 12, e0179588–e0179588, doi:10.1371/journal.pone.0179588 (2017).

41 Langford, D. J. et al. Coding of facial expressions of pain in the laboratory mouse. Nat Methods 7, 447–449, doi:10.1038/nmeth.1455 (2010).

42 Miller, A., Kitson, G., Skalkoyannis, B. & Leach, M. The effect of isoflurane anaesthesia and buprenorphine on the mouse grimace scale and behaviour in CBA and DBA/2 mice. Applied animal behaviour science 172, 58–62, doi:10.1016/j.applanim.2015.08.038 (2015).

43 Jirkof, P., Arras, M. & Cesarovic, N. Tramadol:Paracetamol in drinking water for treatment of post-surgical pain in laboratory mice. Applied Animal Behaviour Science 198, 95–100, doi:10.1016/j.applanim.2017.09.021 (2018).

44 Clark, T. S., Clark, D. D. & Hoyt, R. F., Jr. Pharmacokinetic comparison of sustained-release and standard buprenorphine in mice. Journal of the American Association for Laboratory Animal Science: JAALAS 53, 387–391 (2014).

45 Kendall, L. V. et al. Pharmacokinetics of sustained-release analgesics in mice. J Am Assoc Lab Anim Sci 53, 478–484 (2014).

46 Batka, R. J. et al. The need for speed in rodent locomotion analyses. Anat Rec (Hoboken) 297, 1839–1864, doi:10.1002/ar.22955 (2014).

47 Cendelín, J., Voller, J. & Vožeh, F. Ataxic gait analysis in a mouse model of the olivocerebellar degeneration. Behavioural Brain Research 210, 8–15, doi:https://doi.org/10.1016/j.bbr.2010.01.035 (2010).

48 Jacobs, B. Y., Kloefkorn, H. E. & Allen, K. D. Gait analysis methods for rodent models of osteoarthritis. Curr Pain Headache Rep 18, 456, doi:10.1007/s11916-014-0456-x (2014).

49 Magnusdottir, R. et al. Fracture-induced pain-like behaviours in a femoral fracture mouse model. Osteoporosis International 32, 2347–2359, doi:10.1007/s00198-021-05991-7 (2021).

50 Bizzoca, D. et al. Gait analysis in the postoperative assessment of intertrochanteric femur fractures. J Biol Regul Homeost Agents 34, 345–351. Congress of the Italian Orthopaedic Research Society (2020).

51 Wong, J. et al. Gait patterns after fracture of the femoral shaft in children, managed by external fixation or early hip spica cast. J Pediatr Orthop 24, 463–471, doi:10.1097/00004694-200409000-00003 (2004).

52 Matthiesen, T., Wöhrmann, T., Coogan, T. P. & Uragg, H. The experimental toxicology of tramadol: an overview. Toxicol Lett 95, 63–71, doi:10.1016/s0378-4274(98)00023-x (1998).

53 Dai, X. et al. Gender differences in the antinociceptive effect of tramadol, alone or in combination with gabapentin, in mice. J Biomed Sci 15, 645–651, doi:10.1007/s11373-008-9252-0 (2008).

54 Liu, H. C., Jin, S. M. & Wang, Y. L. Gender-related differences in pharmacokinetics of enantiomers of trans-tramadol and its active metabolite, trans-O-demethyltramadol, in rats. Acta Pharmacol Sin 24, 1265–1269 (2003).

55 Röntgen, V. et al. Fracture healing in mice under controlled rigid and flexible conditions using an adjustable external fixator. J Orthop Res 28, 1456–1462, doi:10.1002/jor.21148 (2010).

56 Fillingim, R. B. & Gear, R. W. Sex differences in opioid analgesia: clinical and experimental findings. Eur J Pain 8, 413–425, doi:10.1016/j.ejpain.2004.01.007 (2004).

57 Smith, J. C. A Review of Strain and Sex Differences in Response to Pain and Analgesia in Mice. Comp Med 69, 490–500, doi:10.30802/AALAS-CM-19-000066 (2019).

58 Patra, D. & Sandell, L. J. Antiangiogenic and anticancer molecules in cartilage. Expert Reviews in Molecular Medicine 14, e10, doi:10.1017/erm.2012.3 (2012).

59 Shukunami, C. & Hiraki, Y. Role of cartilage-derived anti-angiogenic factor, chondromodulin-I, during endochondral bone formation. Osteoarthritis Cartilage 9 Suppl A, S91–101, doi:10.1053/joca.2001.0450 (2001).

60 Lienau, J. et al. Initial vascularization and tissue differentiation are influenced by fixation stability. Journal of Orthopaedic Research 23, 639–645, doi:https://doi.org/10.1016/j.orthres.2004.09.006 (2005).

61 Haffner-Luntzer, M., Fischer, V. & Ignatius, A. Differences in Fracture Healing Between Female and Male C57BL/6J Mice. Front Physiol 12, 712494, doi:10.3389/fphys.2021.712494 (2021).

62 Bove, S. E., Flatters, S. J., Inglis, J. J. & Mantyh, P. W. New advances in musculoskeletal pain. Brain Res Rev 60, 187–201, doi:10.1016/j.brainresrev.2008.12.012 (2009).

63 Cottrell, J. & O’Connor, J. P. Effect of Non-Steroidal Anti-Inflammatory Drugs on Bone Healing. Pharmaceuticals (Basel) 3, 1668–1693, doi:10.3390/ph3051668 (2010).

64 Gerstenfeld, L. C. et al. Differential inhibition of fracture healing by non-selective and cyclooxygenase-2 selective non-steroidal anti-inflammatory drugs. Journal of Orthopaedic Research 21, 670–675, doi:https://doi.org/10.1016/S0736-0266(03)00003-2 (2003).

65 Yassen, A., Olofsen, E., Dahan, A. & Danhof, M. Pharmacokinetic-pharmacodynamic modeling of the antinociceptive effect of buprenorphine and fentanyl in rats: role of receptor equilibration kinetics. J Pharmacol Exp Ther 313, 1136–1149, doi:10.1124/jpet.104.082560 (2005).

66 Yun, M.-H., Jeong, S.-W., Pai, C.-M. & Kim, S.-O. Pharmacokinetic-Pharmacodynamic modeling of the analgesic effect of bupredermTM, in mice. Health 02, 824–831, doi:10.4236/health.2010.28124 (2010).

67 Villiger, J. W. & Taylor, K. M. Buprenorphine: characteristics of binding sites in the rat central nervous system. Life Sci 29, 2699–2708, doi:10.1016/0024-3205(81)90529-4 (1981).

68 Lutfy, K. & Cowan, A. Buprenorphine: a unique drug with complex pharmacology. Curr Neuropharmacol 2, 395–402, doi:10.2174/1570159043359477 (2004).

69 Ohtani, M., Kotaki, H., Sawada, Y. & Iga, T. Comparative analysis of buprenorphine- and norbuprenorphine-induced analgesic effects based on pharmacokinetic-pharmacodynamic modeling. J Pharmacol Exp Ther 272, 505–510 (1995).

70 Lang, A. et al. Collagen I-based scaffolds negatively impact fracture healing in a mouse-osteotomy-model although used routinely in research and clinical application. Acta Biomater 86, 171–184, doi:10.1016/j.actbio.2018.12.043 (2019).

71 Lang, A. et al. MIF does only marginally enhance the pro-regenerative capacities of DFO in a mouse-osteotomy-model of compromised bone healing conditions. Bone 154, 116247, doi:10.1016/j.bone.2021.116247 (2022).

72 Hess, S. E. et al. Home improvement: C57BL/6J mice given more naturalistic nesting materials build better nests. J Am Assoc Lab Anim Sci 47, 25–31 (2008).

73 Miller, A., Kitson, G., Skalkoyannis, B. & Leach, M. The effect of isoflurane anaesthesia and buprenorphine on the mouse grimace scale and behaviour in CBA and DBA/2 mice. Appl Anim Behav Sci 172, 58–62, doi:10.1016/j.applanim.2015.08.038 (2015).

74 Rasband, W. S. ImageJ, <https://imagej.nih.gov/ij/> (1997-2018).

75 Schneider, C. A., Rasband, W. S. & Eliceiri, K. W. NIH Image to ImageJ: 25 years of image analysis. Nature Methods 9, 671–675, doi:10.1038/nmeth.2089 (2012).

76 Noguchi, K., Gel, Y. R., Brunner, E. & Konietschke, F. nparLD: An R Software Package for the Nonparametric Analysis of Longitudinal Data in Factorial Experiments. Journal of Statistical Software 50, 1–23, doi:10.18637/jss.v050.i12 (2012).

77 Kruskal, W. H. & Wallis, W. A. Use of Ranks in One-Criterion Variance Analysis. Journal of the American Statistical Association 47, 583–621, doi:10.1080/01621459.1952.10483441 (1952).

78 Dinno, A. Nonparametric Pairwise Multiple Comparisons in Independent Groups using Dunn’s Test. The Stata Journal 15, 292–300, doi:10.1177/1536867x1501500117 (2015).

79 Dunn, O. J. Multiple comparisons among means. Journal of the American statistical association 56, 52–64 (1961).

