## Supplementary Information for "A Buprenorphine depot formulation provides effective sustained post-surgical analgesia for 72h in mouse femoral fracture models"

Affiliations:

[#]Correspondences:

***Supplementary Methods***

***Health Monitoring – CCR Animal Facility, Charité-Universitätsmedizin Berlin***

Health Monitoring was carried out in accordance with FELASA recommendations.
Species: Mouse
Abbreviations: CULT = Culture; PCR = Polymerase Chain Reaction; IFA = immunofluorescence assay; PATH = Gross Pathology; MICR = Microscopy
POS = positive; NEG = negative; NT = not tested
Housing type: semi-barrier, limited access, protective clothing, IVC

Laboratory: Gesellschaft für innovative Mikroökologie mbH, Waldheimstrasse 47, 14552 Berlin

The following table summarizes the testing results. The study was caried out in between the six months between “latest results” and “previous results”.

| **Test animals** | **Bedding sentinels** | | | |
|---|---|---|---|---|
| | Test frequency | Latest results | Previous results | Testing methods |
| Numbers of animals test | | n = 4 | n = 4 | |
| **Viruses** | | | | |
| Mouse rotavirus (EDIM) | 6 months | NEG | NEG | IFA |
| Mouse hepatitis virus (MHV) | 6 months | NEG | NEG | IFA |
| Murine norovirus (MNV) | 6 months | POS(4) | POS(4) | IFA |
| Mouse parvovirus (MPV) | 6 months | NEG | NEG | IFA |
| Minute virus of mice (MVM) | 6 months | NEG | NEG | IFA |
| Theiler´s murine encephalomyelitis virus (TMEV) | 6 months | NEG | NEG | IFA |
| Lymphocytic choriomeningitis virus (LCMV) | annually | NEG | NT | IFA |
| Mouse adenovirus 1/2 (Fl/K87) | annually | NEG | NT | IFA |
| Mousepox (ectromelia) virus (EV) | annually | NEG | NT | IFA |
| Pneumonia virus of mice (PVM) | annually | NEG | NT | IFA |
| Reovirus type 3 (Reo) | annually | NEG | NT | IFA |
| Sendai virus (SV) | annually | NEG | NT | IFA |
| **Bacteria** | | | | |
| *Rodentibacter spp.* | 6 months | NEG | NEG | CULT |
| *Streptococci spp. ß-haemolytic/not group D* | 6 months | NEG | NEG | CULT |
| *Streptococcus pneumoniae* | 6 months | NEG | NEG | CULT |
| *Helicobacter spp.* | 6 months | POS(2) | POS(2) | PCR |
| *Corynebacterium kutscheri* | annually | NT | NT | CULT |
| *Citrobacter rodentium* | annually | NT | NT | CULT |
| *Salmonella spp.* | annually | NT | NT | CULT |
| *Streptobacillus moniliformis* | annually | NT | NT | PCR |
| *Clostridium piliforme* | annually | NT | NT | IFA |
| *Mycoplasma pulmonis* | annually | NT | NT | IFA |
| **Parasites** | | | | |
| Ectoparasites | 6 months | NEG | NEG | MICR |
| Endoparasites: | 6 months | POS(6) | POS(4) | MICR |
| Aspiculuris spp. | 6 months | NEG | NEG | MICR |
| Syphacia spp. | monthly | NEG | NEG | MICR |
| Chilomastix spp. | 6 months | NEG | NEG | MICR |
| Coccidia spp. | 6 months | NEG | NEG | MICR |
| Entamoeba spp. | 6 months | NEG | NEG | MICR |
| Giardia spp. | 6 months | NEG | NEG | MICR |
| Spironucleus muris | 6 months | NEG | NEG | MICR |
| Trichomonas spp. | 6 months | POS(2) | NEG | MICR |
| other flagellates | 6 months | POS(4) | POS(4) | MICR |
| **Pathological lesions** | 6 months | NEG | NEG | PATH |

***Exclusion of mice and data***

Three male mice were excluded before the beginning of the study due to one sudden death and two urethral obstructions resulting in n = 9 in the following groups: Tramadol rigid, Tramadol flexible and BUP-Depot (Fig. 1A). The following data had to be additionally excluded from analyses: water intake of one cage at 24h after osteotomy due to a leaking bottle (male mice, BUP-Depot, flexible); one male mouse (BUP-Depot, rigid) from composite score and CatWalk analyses, as this animal received an additional injection of Temgesic at 24h post-osteotomy; one female mouse (BUP-Depot, flexible) from stride length analysis (CatWalk) as an technical artifact at 48h post-osteotomy; one female mouse (BUP-Depot, flexible) from limp score analyses at all time points due short-term nerve damage (transient dragging); four samples from ex-vivo *µCT*, histology and immunofluorescence (Tramadol rigid (male), Tramadol flexible (male), Tramadol rigid (female), BUP-Depot flexible (female)) due to insufficient fixation which was not visible until sample collection. In some samples, immunofluorescence staining was not sufficient for analysis and resulted in exclusion of these samples.

### ***Supplementary Results***

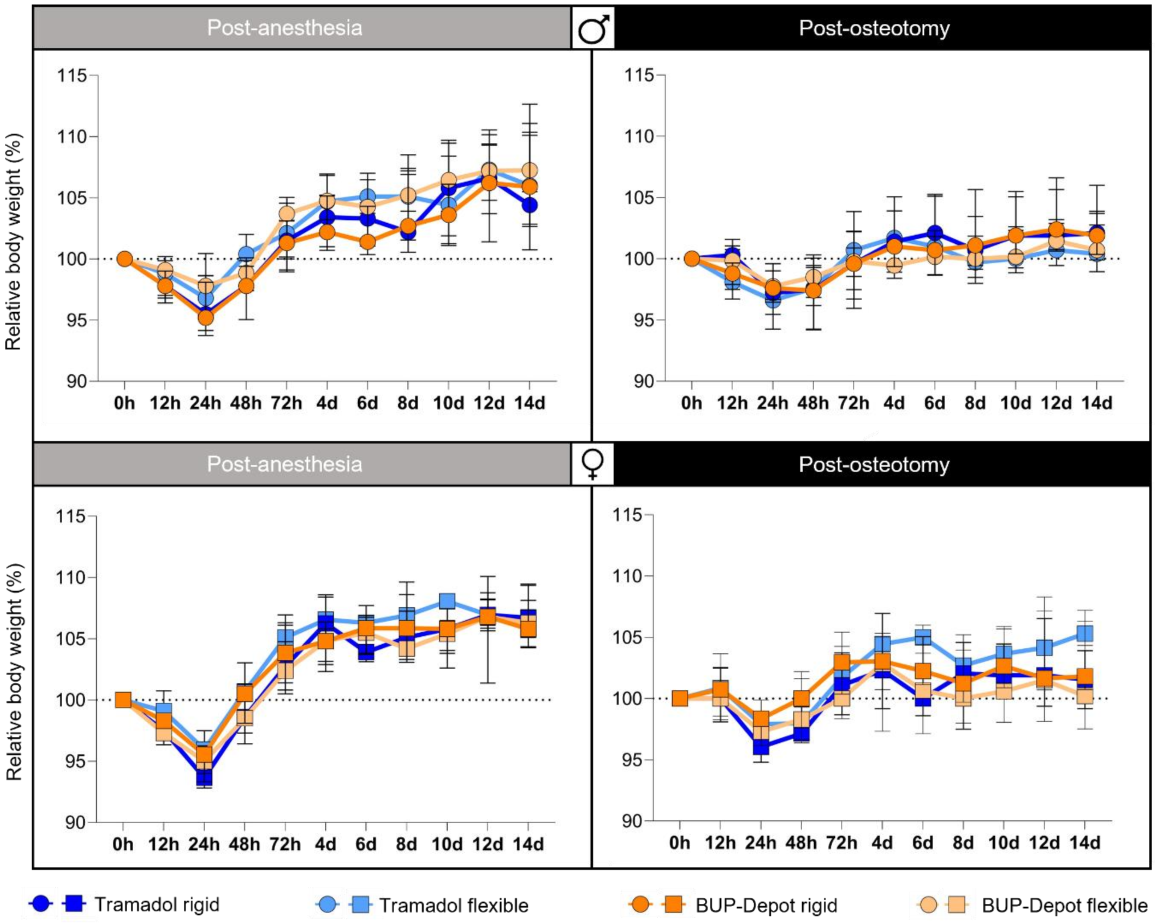


**Figure S1: Relative body weight development over 14 days post-anesthesia and post-osteotomy.** Body weight was measured at 12h, 24h, 48h and 72h post-anesthesia and post-osteotomy and afterwards every other day until osteotomy/euthanasia. Body weight was normalized to the initial value (0h = 100%). All graphs show median with interquartile range for n= 9-10. No statistical analysis was performed.

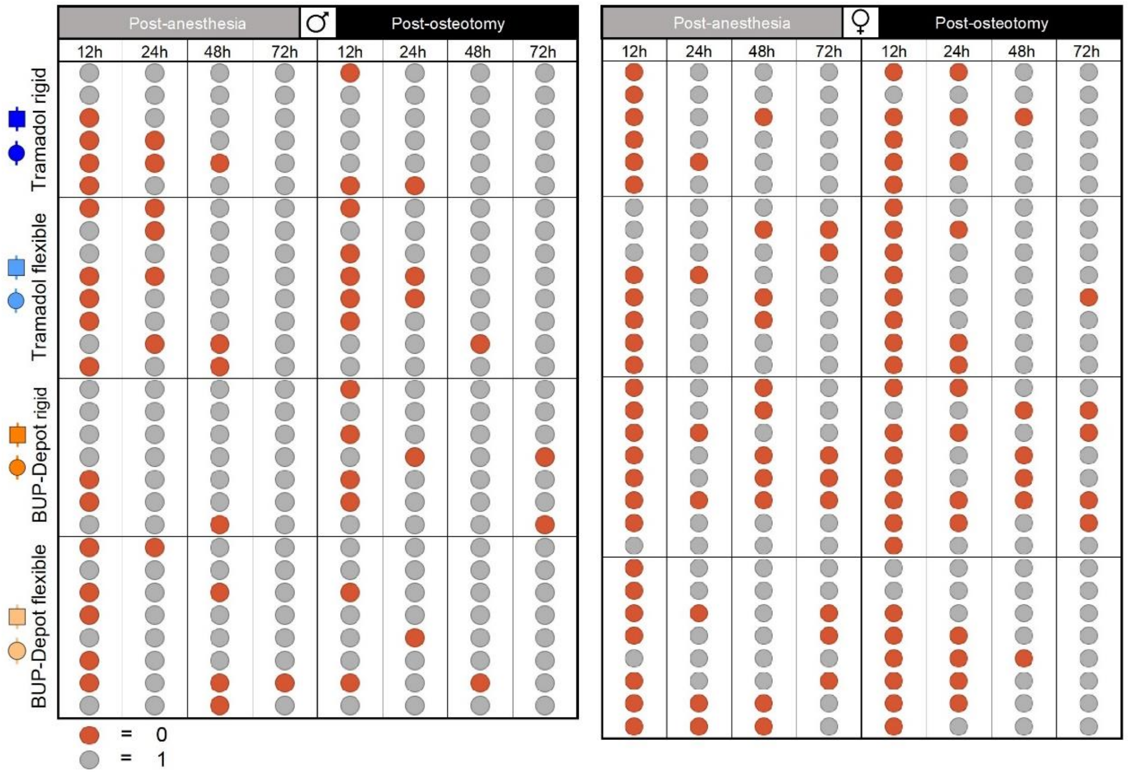


**Figure S2: Defecation during the assessments post-anesthesia and post-osteotomy.** Mice were closely monitored for defecation during the assessments at 12h, 24h, 48h and 72h post-anesthesia and post-osteotomy. n = 6-8.

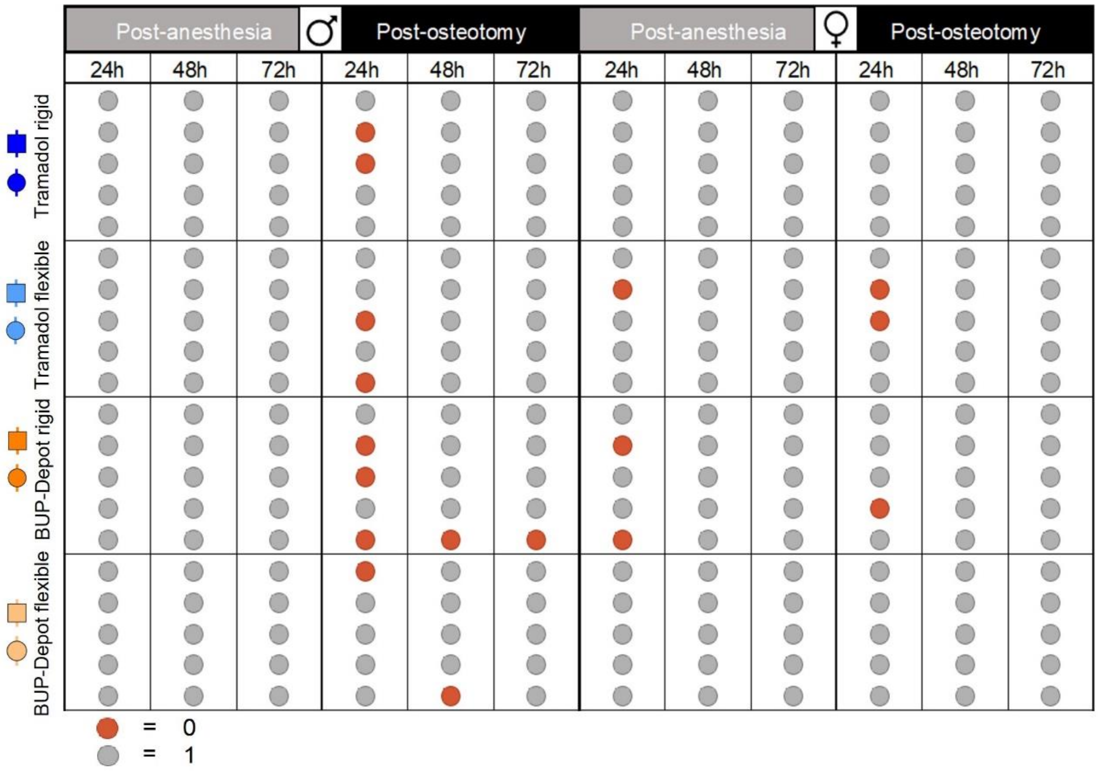


**Figure S3: Explorative behavior post-anesthesia and post-osteotomy. E**xplorative behavior was monitored per cage at 24h, 48h and 72h post-anesthesia and post-osteotomy. n = 5 based on cages (pair housing).

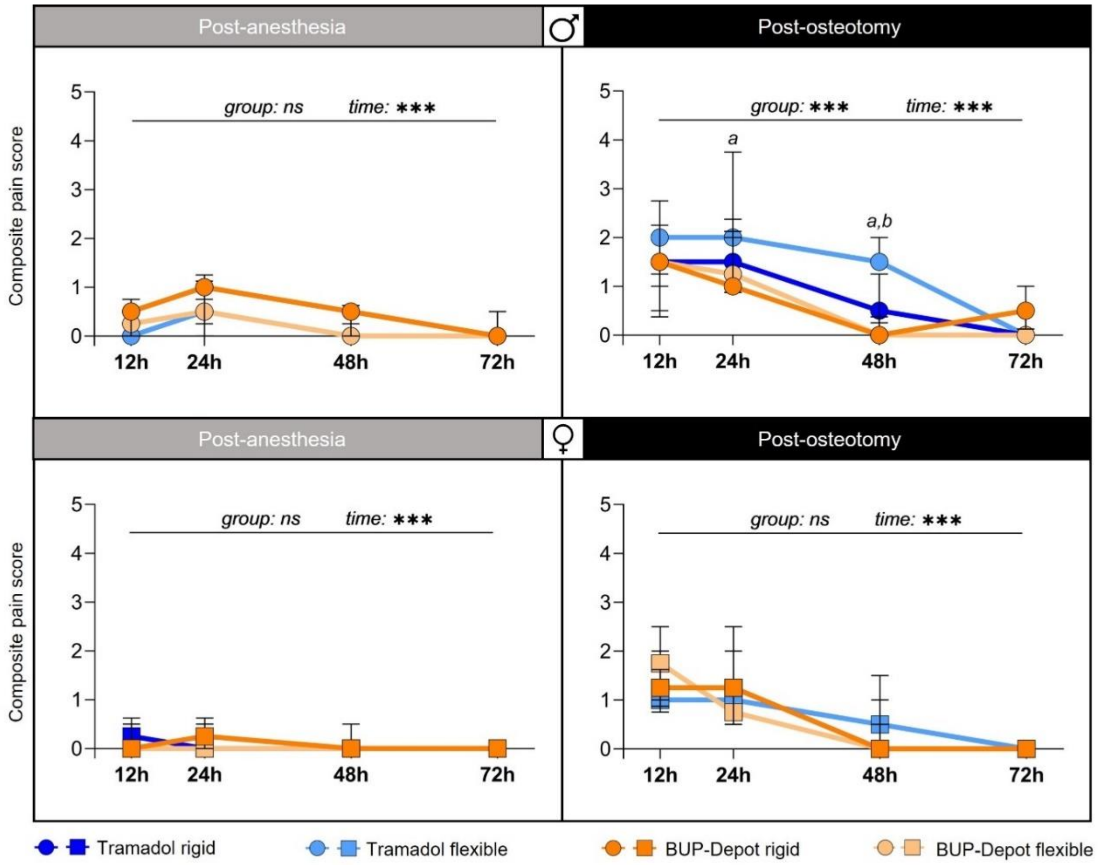


**Figure S4: Composite pain scores post-anesthesia and post-osteotomy.** The composite pain score was measured at 12h, 24h, 48h and 72h post-anesthesia and post-osteotomy. All graphs show median with interquartile range for n= 9-10. Non-parametric ANOVA-type test - main effects of time and of group are represented in the graphs; exact p-values are listed in Supplementary Table S4-S4.3; **p < 0.05, ***p < 0.001*. To determine group differences Dunn's posthoc test with Bonferroni correction was performed. *a* - significant difference Tramadol flexible vs. BUP-Depot flexible; *b* - significant difference Tramadol flexible vs. BUP-Depot rigid.

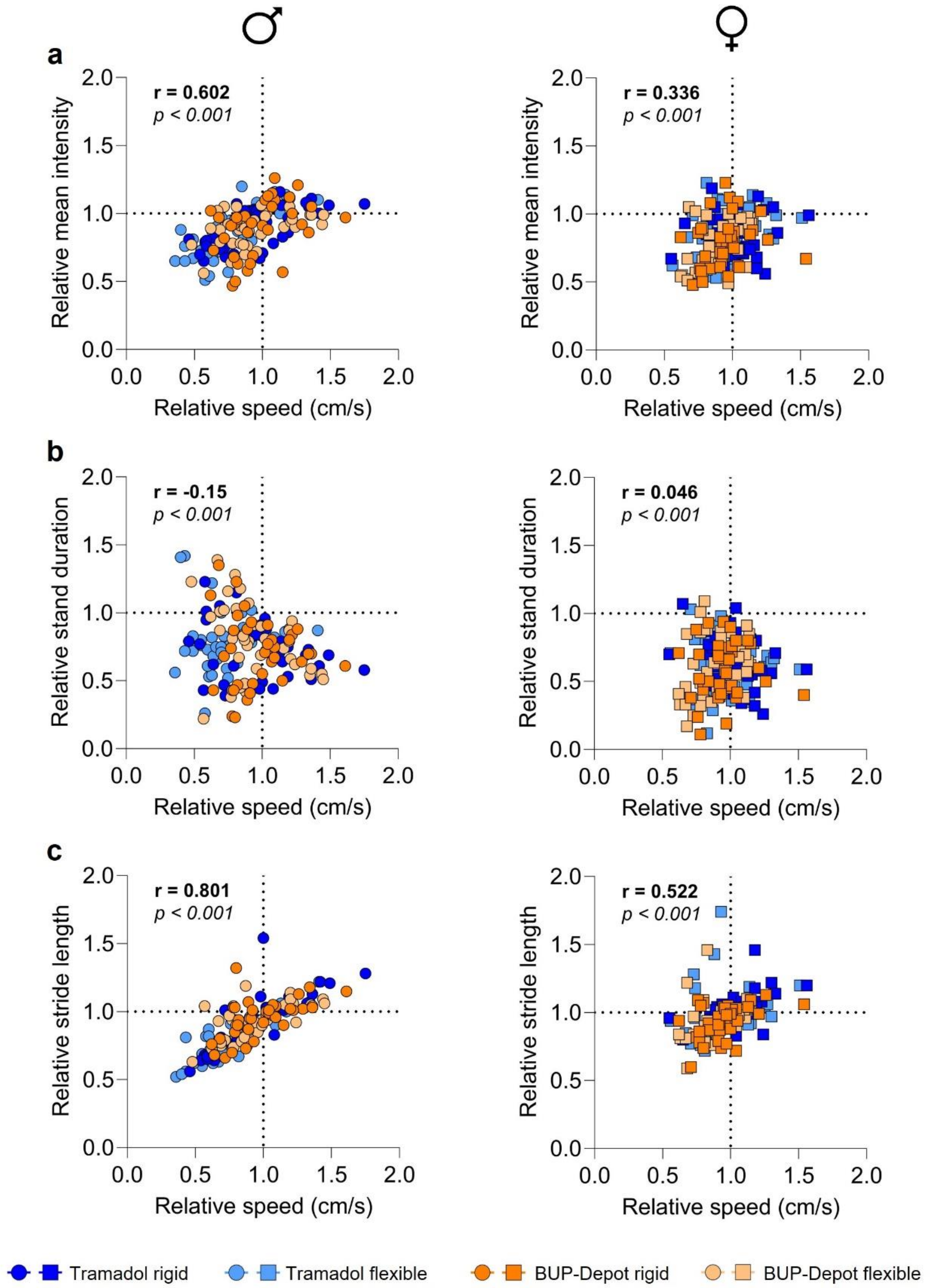


**Figure S5: Spearman correlation between relative velocity and relative mean intensity, relative stand duration and relative stride length.** All parameters were measured at 24h, 48h, 72h and 10d post-osteotomy using the Noldus CatWalk XT. Spearman correlation was carried out combining data from all four timepoints post-osteotomy. Correlation was carried out in Prism (V9).

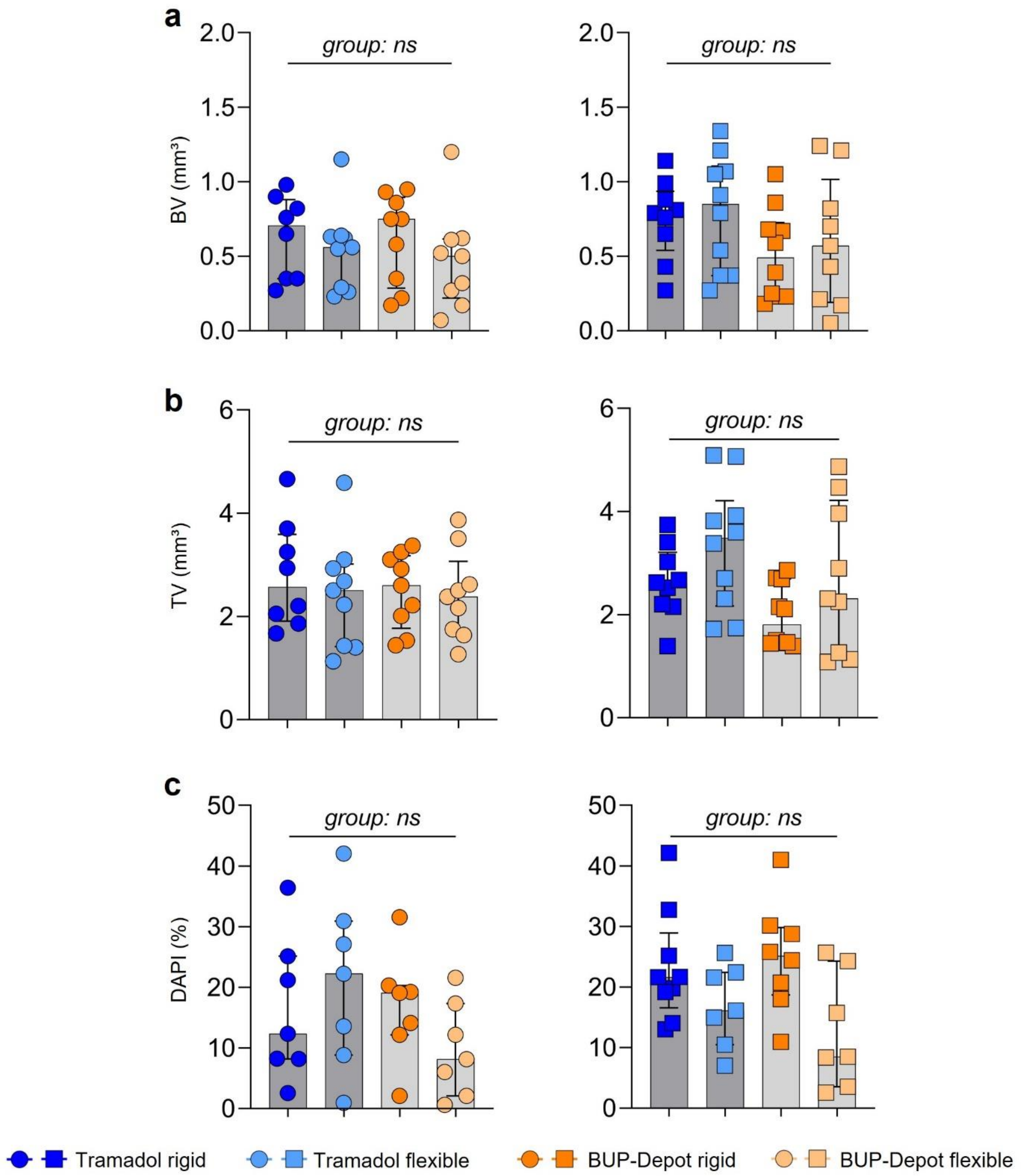


**Figure S6. Analgesic regimes did not negatively affect fracture healing outcome, while the different fixations lead to diverging new bone formation.** (**A**) Bone volume (BV) ($mm^3$), (**B**) total volume (TV) ($mm^3$) and (**C**) DAPI (%) were measured 14d post-osteotomy. All graphs show median with interquartile range for n = 8-10 (**A-B**) and n = 6-9 (**C**). To determine group differences, Kruskal-Wallis, and Dunn's posthoc test with Bonferroni correction was performed in RStudio; exact *p*-values are listed in Supplementary Table **S11-S13.1**.

### *Supplemental Statistics*

**Table S1: *p*-values for body weight**, *nonparametric analysis of longitudinal data (nparLD) (F1-LD-F1 design)*

| | ♂ post-anesthesia | ♂ post-osteotomy | ♀ post-anesthesia | ♀ post-osteotomy |
|---|---|---|---|---|
| Group | 0.631 | 0.887 | **0.0134** | 0.367 |
| Time | **< 0.001** | **< 0.001** | **< 0.001** | **< 0.001** |
| Group:Time | 0.699 | 0.841 | 0.865 | 0.740 |

**Table S1.1: Body weight, females (♀), post-anesthesia**, *Kruskal-Wallis test*

| | *p*-value | chi-squared | df |
|---|---|---|---|
| 12h post-anesthesia | **0.025** | 9.329 | 3 |
| 24h post-anesthesia | 0.051 | 7.784 | 3 |
| 48h post-anesthesia | 0.074 | 6.925 | 3 |
| 72h post-anesthesia | 0.351 | 3.279 | 3 |

**Table S1.2: *p*-values for body weight, females (♀), post-anesthesia,** *Dunns posthoc test (Bonferroni correction), 12h post-anesthesia*

| | ♀, Tramadol flexible | ♀, BUP-Depot rigid | ♀, BUP-Depot flexible |
|---|---|---|---|
| ♀, Tramadol rigid | 0.060 | 0.311 | 1.000 |
| ♀, BUP-Depot rigid | 1.000 | - | |
| ♀, BUP-Depot flexible | **0.031** | 0.186 | |

**Table S1.3: *p*-values for body weight, females (♀), post-anesthesia**, *Dunns posthoc test (Bonferroni correction), 24h post-anesthesia*

| | ♀, Tramadol flexible | ♀, BUP-Depot rigid | ♀, BUP-Depot flexible |
|---|---|---|---|
| ♀, Tramadol rigid | **0.017** | 0.305 | 0.414 |
| ♀, BUP-Depot rigid | 0.765 | - | |
| ♀, BUP-Depot flexible | 0.589 | 1.000 | |

**Table 2: *p*-values for food intake**, *nonparametric analysis of longitudinal data (nparLD) (F1-LD-F1 design)*

| | ♂ post-anesthesia | ♂ post-osteotomy | ♀ post-anesthesia | ♀ post-osteotomy |
|---|---|---|---|---|
| Group | 0.655 | 0.824 | 0.655 | 0.582 |
| Time | **< 0.001** | **< 0.001** | **< 0.001** | **< 0.001** |
| Group:Time | 0.383 | 0.714 | **0.027** | 0.638 |

**Table S3: *p*-values for water intake**, *nonparametric analysis of longitudinal data (nparLD) (F1-LD-F1 design)*

| | ♂ post-anesthesia | ♂ post-osteotomy | ♀ post-anesthesia | ♀ post-osteotomy |
|---|---|---|---|---|
| Group | 0.410 | 0.515 | 0.512 | 0.272 |
| Time | **< 0.001** | **< 0.001** | **< 0.001** | **< 0.001** |
| Group:Time | 0.944 | 0.861 | 0.487 | **0.039** |

**Table S4: *p*-values for the composite pain score**, *nonparametric analysis of longitudinal data (nparLD) (F1-LD-F1 design)*

| | ♂ post-anesthesia | ♂ post-osteotomy | ♀ post-anesthesia | ♀ post-osteotomy |
|---|---|---|---|---|
| Group | 0.184 | **< 0.001** | 0.678 | 0.334 |
| Time | **< 0.001** | **< 0.001** | **< 0.001** | **< 0.001** |
| Group:Time | 0.970 | **0.006** | 0.276 | 0.663 |

**Table S4.1: Composite pain score males** (♂)**, post-osteotomy**, *Kruskal-Wallis test*

| | *p*-value | chi-squared | df |
|---|---|---|---|
| 12h post-osteotomy | 0.182 | 4.861 | 3 |
| 24h post-osteotomy | **0.025** | 9.316 | 3 |
| 48h post-osteotomy | **0.001** | 15.878 | 3 |
| 72h post-osteotomy | 0.234 | 4.271 | 3 |

**Table S4.2: *p*-values for the composite pain score***, Dunns posthoc test (Bonferroni correction), 24h post-osteotomy*

| | ♂, Tramadol flexible | ♂, BUP-Depot rigid | ♂, BUP-Depot flexible |
|---|---|---|---|
| ♂, Tramadol rigid | 0.062 | 1.000 | 1.000 |
| ♂, BUP-Depot rigid | 0.051 | - | |
| ♂, BUP-Depot flexible | **0.020** | 1.000 | |

**Table S4.3: *p*-values for the composite pain score***, Dunns posthoc test (Bonferroni correction), 48h post-osteotomy*

| | ♂, Tramadol flexible | ♂, BUP-Depot rigid | ♂, BUP-Depot flexible |
|---|---|---|---|
| ♂, Tramadol rigid | 0.075 | 0.322 | 1.000 |
| ♂, BUP-Depot rigid | **0.001** | - | |
| ♂, BUP-Depot flexible | **0.009** | 0.941 | |

**Table S5: *p*-values for the delta composite pain score**, *nonparametric analysis of longitudinal data (nparLD) (F1-LD-F1 design)*

| | ♂ post-osteotomy | ♀ post-osteotomy |
|---|---|---|
| Group | **< 0.001** | 0.687 |
| Time | **< 0.001** | **< 0.001** |
| Group:Time | 0.081 | 0.383 |

**Table S5.1: Delta composite pain score, males** (♂)**, post-osteotomy***, Kruskal-Wallis test*

| | *p*-value | chi-squared | df |
|---|---|---|---|
| 12h post-osteotomy | 0.056 | 7.561 | 3 |
| 24h post-osteotomy | **0.031** | 8.909 | 3 |
| 48h post-osteotomy | **< 0.001** | 19.397 | 3 |
| 72h post-osteotomy | 0.278 | 3.847 | 3 |

**Table S5.2: *p*-values for the delta composite pain score**, *Dunns posthoc test (Bonferroni correction), 24h post-osteotomy*

| | ♂, Tramadol flexible | ♂, BUP-Depot rigid | ♂, BUP-Depot flexible |
|---|---|---|---|
| ♂, Tramadol rigid | **0.032** | 1.000 | 1.000 |
| ♂, BUP-Depot rigid | **0.042** | - | |
| ♂, BUP-Depot flexible | 0.077 | 1.000 | |

**Table S5.3: *p*-values for the delta composite pain score**, *Dunns posthoc test (Bonferroni correction), 48h post-osteotomy*

| | ♂, Tramadol flexible | ♂, BUP-Depot rigid | ♂, BUP-Depot flexible |
|---|---|---|---|
| ♂, Tramadol rigid | **0.018** | 0.377 | 1.000 |
| ♂, BUP-Depot rigid | **< 0.001** | - | |
| ♂, BUP-Depot flexible | **0.004** | 0.710 | |

**Table S6: *p*-values for the limp score**, *nonparametric analysis of longitudinal data (nparLD) (F1-LD-F1 design)*

| | ♂ post-osteotomy | ♀ post-osteotomy |
|---|---|---|
| Group | 0.436 | 0.108 |
| Time | **< 0.001** | **0.018** |
| Group:Time | 0.600 | 0.612 |

**Table S7: *p*-values for CatWalk - relative velocity**, *nonparametric analysis of longitudinal data (nparLD) (F1-LD-F1 design)*

| | ♂ post-anesthesia | ♂ post-osteotomy | ♀ post-anesthesia | ♀ post-osteotomy |
|---|---|---|---|---|
| Group | 0.266 | **0.034** | 0.563 | **0.035** |
| Time | **0.003** | **< 0.001** | **0.007** | **< 0.001** |
| Group:Time | 0.662 | 0.133 | 0.530 | 0.092 |

**Table S7.1: CatWalk - relative velocity, males (♂), post-osteotomy**, *Kruskal-Wallis test*

| | *p*-value | chi-squared | df |
|---|---|---|---|
| 24h post-osteotomy | **0.052** | 7.735 | 3 |
| 48h post-osteotomy | **0.038** | 8.428 | 3 |
| 72h post-osteotomy | 0.097 | 6.316 | 3 |
| 10d post-osteotomy | 0.110 | 6.035 | 3 |

**Table S7.2: *p*-values for CatWalk - relative velocity, males (♂), post-osteotomy**, *Dunns posthoc test (Bonferroni correction), 24h post-osteotomy*

| | ♂, Tramadol flexible | ♂, BUP-Depot rigid | ♂, BUP-Depot flexible |
|---|---|---|---|
| ♂, Tramadol rigid | 0.119 | 1.000 | 1.000 |
| ♂, BUP-Depot rigid | **0.034** | - | |
| ♂, BUP-Depot flexible | 0.101 | 1.000 | |

**Table S7.3: *p*-values for CatWalk - relative velocity, males (♂), post-osteotomy**, *Dunns posthoc test (Bonferroni correction), 48h post-osteotomy*

| | ♂, Tramadol flexible | ♂, BUP-Depot rigid | ♂, BUP-Depot flexible |
|---|---|---|---|
| ♂, Tramadol rigid | 0.053 | 1.000 | 1.000 |
| ♂, BUP-Depot rigid | **0.029** | - | |
| ♂, BUP-Depot flexible | 0.157 | 1.000 | |

**Table S7.4: CatWalk - relative velocity, females (♀), post-osteotomy**, *Kruskal-Wallis test*

| | *p*-value | chi-squared | df |
|---|---|---|---|
| 24h post-osteotomy | **0.009** | 11.470 | 3 |
| 48h post-osteotomy | 0.259 | 4.026 | 3 |
| 72h post-osteotomy | 0.180 | 4.894 | 3 |
| 10d post-osteotomy | 0.166 | 5.088 | 3 |

**Table S7.5: *p*-values for CatWalk - relative velocity, females (♀), post-osteotomy**, *Dunns posthoc test (Bonferroni correction), 24h post-osteotomy*

| | ♀, Tramadol flexible | ♀, BUP-Depot rigid | ♀, BUP-Depot flexible |
|---|---|---|---|
| ♀, Tramadol rigid | 0.074 | 0.162 | **0.003** |
| ♀, BUP-Depot rigid | 1.000 | - | |
| ♀, BUP-Depot flexible | 0.855 | 0.583 | |

**Table S8: *p*-values for CatWalk - relative mean intensity**, *nonparametric analysis of longitudinal data (nparLD) (F1-LD-F1 design)*

| | ♂ post-osteotomy | ♀ post-osteotomy |
|---|---|---|
| Group | 0.489 | 0.653 |
| Time | **< 0.001** | **< 0.001** |
| Group:Time | 0.736 | 0.096 |

**Table S9: *p*-values for CatWalk - relative stand duration**, *nonparametric analysis of longitudinal data (nparLD) (F1-LD-F1 design)*

| | ♂ post-osteotomy | ♀ post-osteotomy |
|---|---|---|
| Group | 0.821 | 0.926 |
| Time | **< 0.001** | **0.005** |
| Group:Time | 0.692 | 0.468 |

**Table S10: *p*-values for CatWalk - relative stride length**, *nonparametric analysis of longitudinal data (nparLD) (F1-LD-F1 design)*

| | ♂ post-osteotomy | ♀ post-osteotomy |
|---|---|---|
| Group | **0.024** | 0.328 |
| Time | **< 0.001** | 0.053 |
| Group:Time | 0.175 | 0.580 |

**Table S10.1: CatWalk - relative stride length, males (♂), post-osteotomy**, *Kruskal-Wallis test*

| | *p*-value | chi-squared | df |
|---|---|---|---|
| 24h post-osteotomy | **0.044** | 8.084 | 3 |
| 48h post-osteotomy | 0.119 | 5.845 | 3 |
| 72h post-osteotomy | 0.112 | 5.987 | 3 |
| 10d post-osteotomy | 0.055 | 7.595 | 3 |

**Table S10.2: *p*-values for CatWalk - relative stride length, males** (♂), *Dunns posthoc test (Bonferroni correction), 24h post-osteotomy*

| | ♂, Tramadol flexible | ♂, BUP-Depot rigid | ♂, BUP-Depot flexible |
|---|---|---|---|
| ♂, Tramadol rigid | 0.606 | 0.423 | 1.000 |
| ♂, BUP-Depot rigid | **0.018** | - | |
| ♂, BUP-Depot flexible | 0.151 | 1.000 | |

**Table S11: *Ex-vivo* µCT: BV/TV, BV, TV**, *Kruskal-Wallis test*

| | ♂ | | | ♀ | | |
|---|---|---|---|---|---|---|
| | *p*-value | chi-squared | df | *p*-value | chi-squared | df |
| BV/TV | 0.947 | 0.369 | 3 | 0.263 | 3.983 | 3 |
| BV ($mm^3$) | 0.400 | 2.947 | 3 | 0.248 | 4.124 | 3 |
| TV ($mm^3$) | 0.848 | 0.806 | 3 | 0.081 | 6.728 | 3 |

**Table S12: Histology (Movat's Pentachrome)**, *Kruskal-Wallis test*

| | ♂ | | | ♀ | | |
|---|---|---|---|---|---|---|
| | *p*-value | chi-squared | df | *p*-value | chi-squared | df |
| Relative bone fraction (%) | 0.907 | 0.556 | 3 | 0.089 | 6.521 | 3 |
| Relative cartilage fraction (%) | 0.375 | 3.109 | 3 | 0.165 | 5.094 | 3 |

**Table S13: Immunofluorescence**, *Kruskal-Wallis test*

| | ♂ | | | ♀ | | |
|---|---|---|---|---|---|---|
| | *p*-value | chi-squared | df | *p*-value | chi-squared | df |
| Endomucin (%) | 0.241 | 6.728 | 3 | **0.045** | 8.066 | 3 |
| DAPI (%) | 0.281 | 3.823 | 3 | 0.069 | 7.079 | 3 |

**Table S13.1: Endomucin (%), females** (♀), *Dunns posthoc test (Bonferroni correction)*

| | ♀, Tramadol flexible | ♀, BUP-Depot rigid | ♀, BUP-Depot flexible |
|---|---|---|---|
| ♀, Tramadol rigid | 0.100 | 1.000 | 0.223 |
| ♀, BUP-Depot rigid | 0.083 | - | |
| ♀, BUP-Depot flexible | 1.000 | 0.186 | |
